# BLOS1 mediates kinesin switch during endosomal recycling of LDL receptor

**DOI:** 10.1101/2020.04.22.056135

**Authors:** Chang Zhang, Chanjuan Hao, Guanghou Shui, Wei Li

## Abstract

Low-density lipoprotein receptor (LDLR) in hepatocytes plays a key role in normal clearance of circulating LDL and in whole body cholesterol homeostasis. The trafficking of LDLR is highly regulated in clathrin-dependent endocytosis, endosomal recycling and lysosomal degradation. Current studies have been focusing on its endocytosis and degradation. However, the detailed molecular and cellular mechanisms underlying its endosomal recycling are largely unknown. We found that BLOS1, a shared subunit of BLOC-1 and BORC, is involved in LDLR endosomal recycling. Loss of BLOS1 leads to less membrane LDLR and impairs LDL clearance from plasma in hepatocyte-specific BLOS1 knockout mice. BLOS1 interacts with kinesin-3, and that BLOS1 acts as a new adaptor for kinesin-2 to coordinate kinesin-3 and kinesin-2 during the long-range transport of recycling endosomes (REs) to plasma membrane along microtubule tracks to overcome hurdles at microtubule intersections. These findings provide new insights into RE’s anterograde transport and the pathogenesis of dyslipidemia.

## INTRODUCTION

Low-density lipoprotein receptor (LDLR) is essential for cellular uptake of cholesterol-carrying low-density lipoproteins (LDL) and plays a crucial role in cholesterol homeostasis in mammals (Goldstein & Brown, 2009, 2015). After the binding of LDL on the cell surface, LDLR enters early endosomes through clathrin-mediated endocytosis (CME) and is further recycled to the cell membrane by the RAB4-dependent fast recycling pathway or the RAB11A-dependent slow recycling pathway (Ullrich, Reinsch, Urbé, Zerial, & Parton, 1996; van der Sluijs et al., 1992; Wijers, Kuivenhoven, & van de Sluis, 2015). In the slow recycling pathway, receptors are first sorted into the perinuclear endocytic recycling compartment (ERC) and then transported to the cell periphery by recycling endosomes (REs) (Naslavsky & Caplan, 2018; Yamashiro, Tycko, Fluss, & Maxfield, 1984). Due to the long distance in trafficking, the slow recycling pathway is subjected to complex regulation. Factors including RAB11A effectors (RAB11-FIPs) and Eps15-homology domain-containing proteins (EHDs) participate in the trafficking from early endosome (EE) to the ERC (Horgan & McCaffrey, 2009; Naslavsky & Caplan, 2011).

N-kinesins, the motor proteins that drive anterograde transport on microtubules, have also been reported to function in slow recycling (Hirokawa, Noda, Tanaka, & Niwa, 2009). A kinesin-3 member KIF13A cooperates with RAB11A to generate and drive the peripheral transport of RE tubules (Delevoye et al., 2014; Nakagawa et al., 2000). In addition, KIF13A is essential for cargo delivery from RE to maturing melanosomes in melanocytes (Delevoye et al., 2009). KIF16B, another member of kinesin-3, functions in regulating the motility of early endosomes and degradation of EGF receptor, and in the transcytosis of TfR in polarized epithelial cells (Hoepfner et al., 2005; Perez Bay et al., 2013). A kinesin-2 member KIF3B has also been implicated in the recycling of TfR through the interaction with RAB11-FIP5 (Schonteich et al., 2008). Whether and how these different kinesins coordinate in the transport of REs are largely unknown.

BLOS1 is a shared subunit of BLOC-1 (biogenesis of lysosome-related organelles complex-1) and BORC (BLOC-one-related complex) (Scott, Wang, Wu, Thapa, & Sack, 2018). BLOC-1 is required for the biogenesis of cell type specific lysosome-related organelles, such as melanosomes in melanocytes and dense granules in platelets (Bowman, Bi-Karchin, Le, & Marks, 2019; Wei & Li, 2013). BORC recruits ARL8 and couples lysosomes to kinesins for anterograde transport on specific microtubule tracks, and mediates the trafficking of synaptic vesicle precursors in neurons (Guardia, Farías, Jia, Pu, & Bonifacino, 2016; Niwa et al., 2017). There is increasing evidence that BLOC-1 is involved in the formation of endosomal tubular structures through cooperation with microtubule- and actin-associated machineries (Delevoye et al., 2016; Ryder et al., 2013).

Here we show that BLOS1 regulates LDLR recycling in hepatocytes. BLOS1 acts as an adaptor of kinesin-2 and coordinates kinesin-2 and kinesin-3 in the anterograde transport of REs. Dysfunction of kinesin-2 or BLOS1 results in impaired trafficking of REs and KIF13A-positive tubular structures characterized by impassability at specific microtubule-microtubule intersections. Consequently, liver-specific *Bloc1s1* knockout mice exhibit reduced LDLR in the liver and elevated plasma LDL levels due to alternative lysosome degradation of LDLR after impaired endocytic recycling.

## RESULTS

### Liver-specific knockout of *Bloc1s1* in mice leads to abnormal lipid metabolism

Previous studies have shown that constitutive knockout of *Bloc1s1* in mice produces embryonic lethality (Scott et al., 2014; Zhang et al., 2014). To study the function of BLOS1 in liver lipid metabolism, we generated a conditional knockout mouse mutant (cKO mice) by crossing *Bloc1s1* floxed mice (loxp mice) with Albumin (Alb)-cre mice, while littermates lacking *cre* gene served as control group (**Fig. S1a**). Deletion of *Bloc1s1* was confirmed in hepatocytes isolated from cKO mouse liver by genomic PCR (**Fig. S1b**). Because antibody to BLOS1 was not applicable, we used two other BLOC-1 subunits (Pallidin and Dysbindin) to monitor the loss of BLOS1 as the depletion of BLOS1 leads to the destabilization of other BLOC-1 subunit (Zhang et al., 2014). Indeed, the content of both subunits was significantly reduced in cKO mouse livers and hepatocytes, suggesting the depletion of BLOS1 (**Fig. 1a,b**).

**Figure 1.**
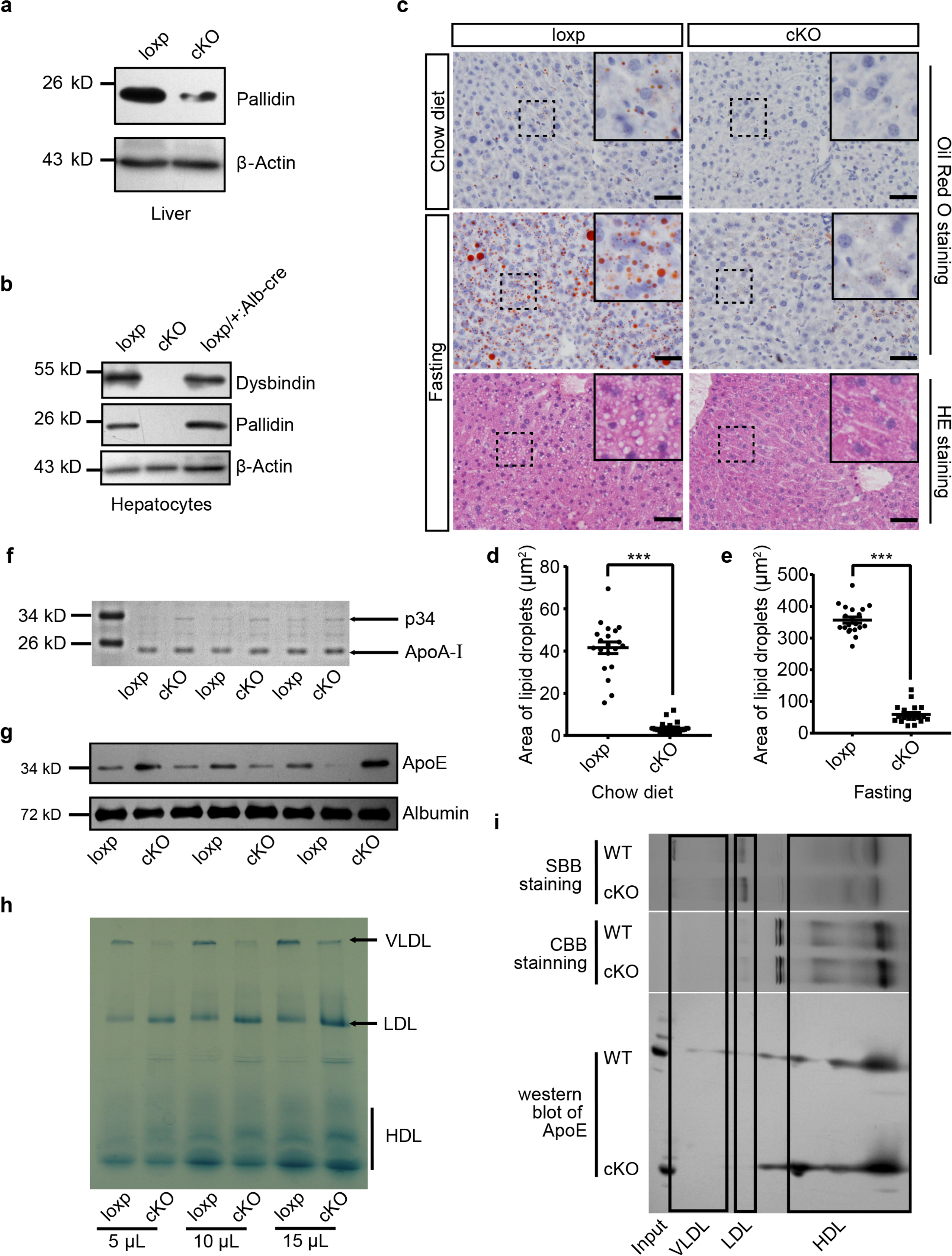
Mice with hepatocyte-specific deletion of BLOS1 showed abnormal lipid metabolism. (**a, b**) Immunoblot of the destabilized BLOC-1 subunits (Pallidin and Dysbindin) in lysates of liver (a) and purified primary hepatocytes (b) from loxp and cKO mice. The loxp heterozygous mice in (b) were used as another control. Note that the weak band of Pallidin in (a) indicates the protein from other non-hepatocytes in the liver. (**c**) Representative images showing the lipid droplets in liver frozen sections of loxp and cKO mice under indicated conditions. For mice after fasting, both Oil Red O staining and HE staining results are displayed. Magnified insets of boxed areas are placed on the top right corners of each picture. Scale bars, 50 μm. (**d, e**) Average area of lipid droplets in 20 random 50 μm × 50 μm square regions before (d) and after (e) starvation showing reduced lipid droplet content in cKO mice. Quantifications were performed on Oil Red O stained sections. Mean ± SEM. Two-tailed t test, ***p < 0.001. (**f**) Coomassie brilliant blue staining (CBB staining) of plasma proteins in loxp and cKO mice after separation by SDS-PAGE. (**g**) Immunoblot of ApoE in plasma of different loxp and cKO mice, albumin is a loading control. (**h**) Lipoproteins prestained by Sudan Black B in plasma of loxp and cKO mice were separated by 4-15% gradient native PAGE at different loading volume of plasma. (**i**) Immunoblot of ApoE in lipoproteins separated by native PAGE and a second dimensional SDS-PAGE. Gel slices of prestained lipoproteins and CBB stained proteins were used to determine the location of different lipoproteins in immunoblots. SBB, Sudan Black B. See also **Fig. S1**.

To investigate whether BLOS1 deficiency could affect liver lipid droplet content, we performed Oil Red O staining on frozen sections of mouse livers and found that cKO mice had fewer lipid droplets in the liver when fed on chow diet, and the accumulation of lipid droplets after starvation was largely inhibited in cKO mouse liver (**Fig. 1c-e**). Considering that many important constituents of plasma are secreted by liver, we examined plasma samples of cKO mice and their control littermates by non-reduced SDS-PAGE analysis. Although there were no changes in several abundant plasma proteins, such as albumin and transferrin, a protein band at about 34 kD was consistently increased in plasma samples from cKO mice (**Fig. 1f**). Mass spectrometry identified this protein as apolipoprotein E (ApoE) (**Fig. S1c**), and immunoblotting confirmed the increase of ApoE in cKO mice plasma (**Fig. 1g**).

ApoE is a core protein component of very-low-density lipoprotein (VLDL) and high-density lipoprotein (HDL). For this reason, lipoproteins in plasma samples from cKO and loxp mice were prestained with Sudan Black B and then separated by gradient native polyacrylamide gel electrophoresis. Using lipoproteins purified from pooled mouse plasma by sequential ultracentrifugation, we determined different lipoprotein bands in native gels (**Fig. S1d**, left). We observed that in cKO mouse plasma, VLDL level was decreased, low-density lipoprotein (LDL) was increased, and HDL had a tendency to increase (**Fig. 1h**). Native gels subjected to Oil Red O staining also showed similar results (**Fig. S1d**, right). The reduction of VLDL in cKO plasma apparently could not account for the increase of ApoE content. We wondered whether ApoE is increased in the HDL fraction of lipoproteins. To address this point, we separated unstained lipoproteins from cKO and loxp mice using native gels, and then performed a second dimensional SDS-PAGE and immunoblotting to the gel slices of target lanes containing separated lipoproteins. Indeed, we observed that there was more ApoE in the HDL slice of cKO mice (**Fig. 1i**).

Collectively, these results suggest that cKO mice have abnormal lipid metabolism, mainly characterized by reduced lipid droplets in the liver, and disordered plasma lipoprotein compositions with increase of LDL, reduction of VLDL, and increase of ApoE in HDL.

### BLOS1 deficiency reduces membrane LDLR in hepatocytes

Despite the reduced VLDL content, a common constituent of LDL and VLDL, ApoB, was significantly increased in cKO mouse plasma (**Fig. S2a**), suggesting that increase of LDL in cKO mouse plasma was likely the main effect of BLOS1 deficiency on lipoproteins. VLDL is largely converted to LDL in plasma. The reduced VLDL in cKO mouse plasma is unlikely the major cause of increased LDL. Liver is the primary site for plasma LDL clearance, the accumulation of LDL in cKO mouse plasma suggested that LDL clearance by cKO mouse liver could be impaired. To test this hypothesis, we examined the endocytosis of LDL in primary hepatocytes using purified mouse LDL labeled with DiI (LDL-DiI). We observed that LDL endocytosis in cKO hepatocytes was reduced at the initial stage (**Fig. 2a, b**), suggesting that less LDL binds to the cell membrane of cKO hepatocytes. We then detected the LDLR protein level by immunoblotting. We found a significant reduction of LDLR in cKO mouse liver (**Fig. 2c, f**; and **Fig. S2b**).

**Figure 2.**
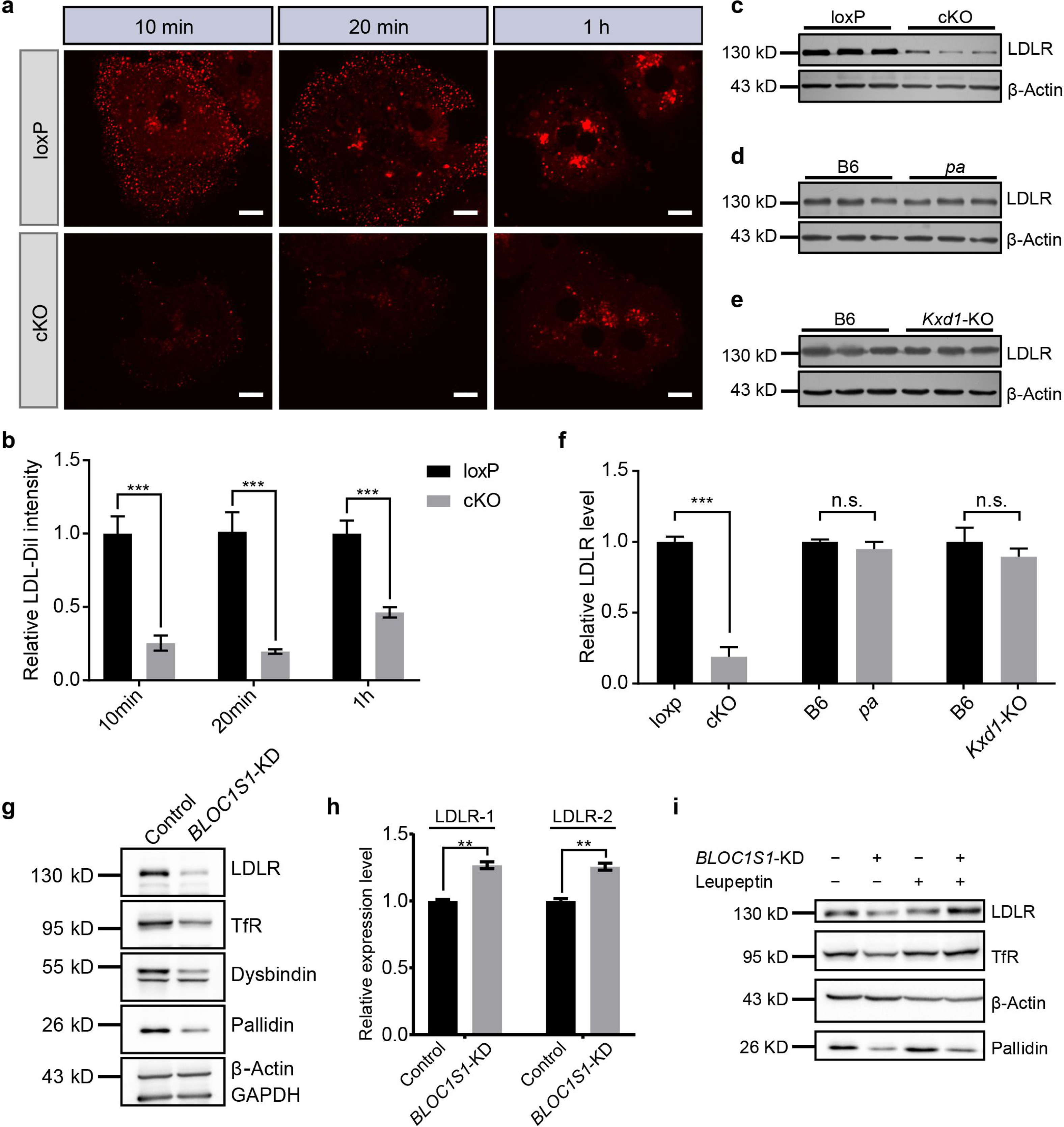
BLOS1 regulates LDLR membrane trafficking and interacts with LDLR. (**a**) Endocytosis of DiI labeled LDL in hepatocytes from loxp and cKO mice after indicated time period. Scale bars, 10 μm. (**b**) Average intensity of LDL-DiI signal after endocytosis in loxp (n = 10 [10min], n = 10 [20min], n = 11 [2h]) and cKO (n = 8 [10min], n = 9 [20min], n = 8 [2h]) hepatocytes at indicated time points. Mean ± SEM. Two-tailed t test, ***p < 0.001. (**c-e**) Immunoblot of LDLR in cKO, pa and Kxd1-KO mouse livers showing the decreased LDLR in cKO mice, but not in pa or Kxd1-KO mice. (**f**) Quantification of relative LDLR protein level (normalized to β-Actin) in pa (n =3), Kxd1-KO (n = 6) and cKO (n =5) mice and their control goups. Mean ± SEM. Two-tailed t test; n.s., not significant; ***p < 0.001. (**g**) Detection of LDLR and TfR in BLOC1S1 stable knockdown Hep G2 cells (Bloc1s1-KD cells). Destabilization of Pallidin or Dysbindin is also shown. (**h**) qRT-PCR of LDLR normalized to GAPDH using two different primer pairs (LDLR-1 and LDLR-2) in control and BLOC1S1-KD cells. Mean ± SEM, three repeats. Two-tailed t test, **p < 0.01. (**i**) Recovery of LDLR level in Bloc1s1-KD cells after the inhibition of lysosomal degradation by treating with leupeptin. See also **Fig. S2**.

The reduction of LDLR in cKO mouse liver was not attributable to the dysfunction of either the BLOC-1 or BORC complexes, as both the *pa* mice (lack of Pallidin subunit in BLOC-1) and *Kxd1*-KO mice (lack of KXD1 subunit in BORC) showed no obvious changes in liver LDLR levels (**Fig. 2d-f**). This suggests that the effect on LDLR is specific for BLOS1 alone. Considering that the uptake of LDL by other tissues (except the liver) is unaffected in cKO mice, the increase of plasma LDL is mainly attributable to impaired liver clearance of LDL.

The reduction of LDL caused by BLOS1 deficiency was further confirmed in Hep G2 cells with stable knockdown of *BLOC1S1* (*BLOC1S1*-KD cells) (**Fig. 2g**). The potential down-regulation effect of BLOS1 deficiency on LDLR transcriptional levels was excluded by qRT-PCR in two different sets of primers (**Fig. 2h**) (actually, the mRNA level of LDLR was slightly increased in *BLOC1S1*-KD cells), suggesting a post-transcriptional effect on LDLR. In addition, reduction of LDLR level in *BLOC1S1*-KD cells was recovered after the inhibition of lysosomal degradation by leupeptin (**Fig. 2i**), suggesting excessive degradation of LDLR upon the loss of BLOS1. We also noted that the content of PCSK9, a known negative regulator of LDLR degradation, was not affected in cKO mouse livers (**Fig. S2b**).

We next investigated whether BLOS1 could directly interact with LDLR. LDLR co-immunoprecipitated with FLAG-tagged BLOS1 (**Fig. S2c**) and GST-BLOS1 fusion protein could pull down LDLR from liver tissue lysate (**Fig. S2d**), indicating an interaction between BLOS1 and LDLR. We further constructed different truncations of BLOS1, and then performed the GST pull-down experiments using mouse liver lysates. The interacting region on BLOS1 was narrowed down to residues 76-100 (**Fig. S2e-g, i**). Furthermore, the cytosolic domain of LDLR at its C-terminus could pull down BLOS1 (**Fig. S2h, i**).

Taken together, these results showed that loss of BLOS1 in liver could lead to impaired LDL clearance which is caused by excessive lysosomal degradation of LDLR, and residues 76-100 on BLOS1 interacts with the cytosolic domain of LDLR at the C-terminus.

### BLOS1 localizes to microtubules and interacts with kinesins

To further explore the underlying mechanism of how BLOS1 regulates LDLR level, we first performed immunocytochemistry (ICC) to observe the subcellular distribution of BLOS1. In Hep G2 cells, overexpressed BLOS1 with different tags all showed puncta distribution patterns in the cytosol (**Fig. 3a**). When fused to a large tag such as GFP, BLOS1-GFP mostly formed aggregates (**Fig. 3a**, left). However, when fused with small tags such as Myc, FLAG or HA, BLOS1 localized to smaller puncta structures distributed more evenly in the cytosol (**Fig. 3a**, right). We used small tagged BLOS1 in the following immunostaining assays in Hep G2 cells. We co-stained BLOS1 overexpressing cells with different organelle markers and found that almost no colocalization of BLOS1 with mitochondria (labeled by Cytochrome C) was observed (**Fig. 3b**), but BLOS1 partially colocalized with lysosome/MVB marker CD63 (**Fig. 3c**). In addition, a small fraction of BLOS1 colocalized with the early endosome marker EEA1 (**Fig. 3d**), and more BLOS1 puncta colocalized with TfR labeled recycling endosomes (REs) (**Fig. 3e**) or LDLR positive vesicles (**Fig. 3f**, see also **Fig. 3g** for quantification), indicating a potential role of BLOS1 in the endocytic and recycling pathway.

**Figure 3.**
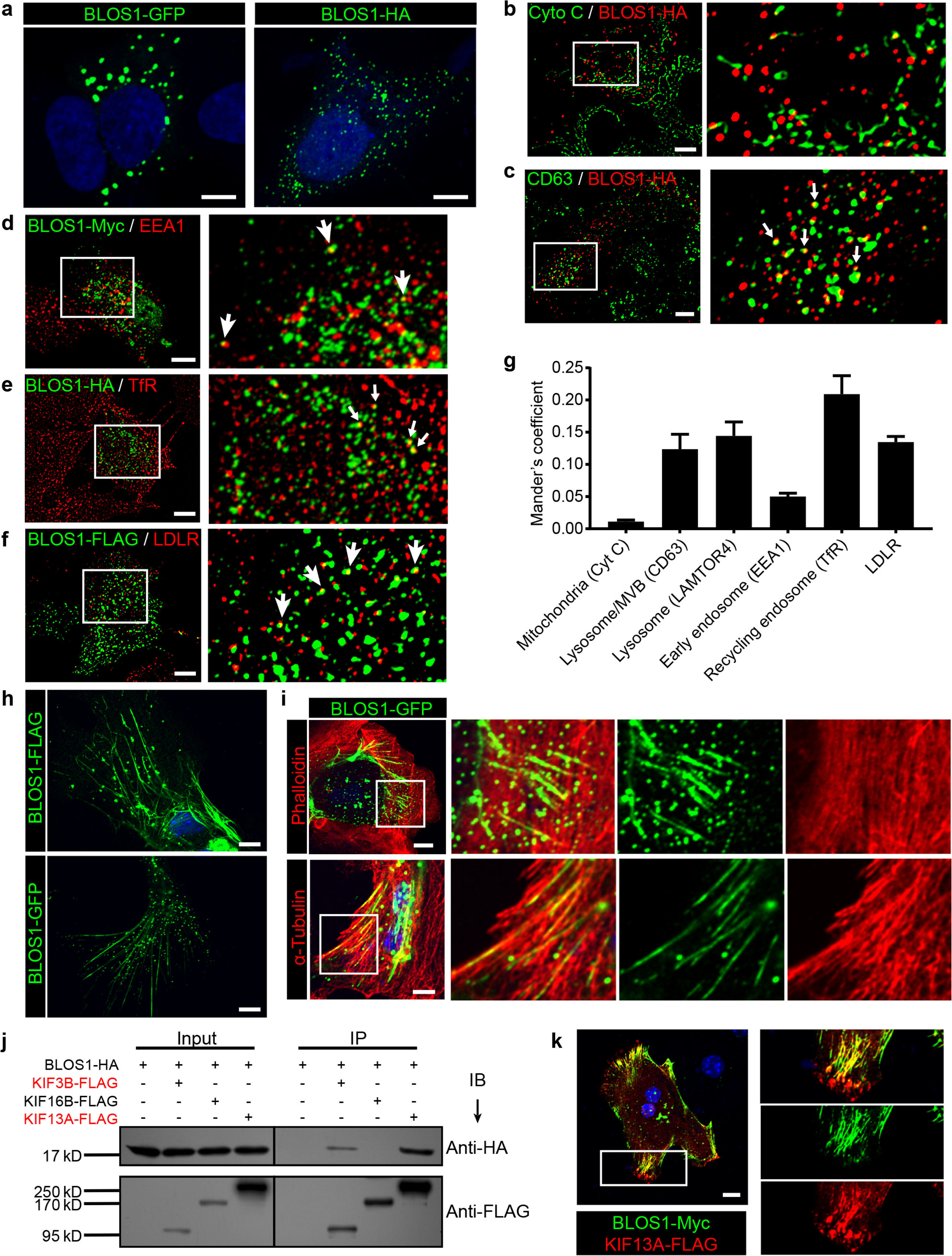
BLOS1 localizes to microtubules and interacts with kinesins. (**a**) Representative confocal images of puncta patterns of BLOS1-GFP-C2, BLOS1-GFP-N2 and immunofluorescence stained BLOS1-Myc and BLOS1-HA in Hep G2 cells. (**b-f**) BLOS1-HA partially colocalizes with multivesicular body/lysosome marker CD63 (c, white arrows), early endosome marker EEA1 (d), recycling endosome marker TfR (e) and LDLR vesicles (f) in Hep G2 cells, while almost no colocalization was observed between BLOS1-HA and mitochondria marker Cytochrome C (b). Magnified insets of boxed areas are shown on right. (**g**) Quantification of Mander’s colocalization coefficient showing the percentage of BLOS1 that colocalizes with other proteins or organelle markers. n = 5, 6, 6, 5, 6 and 6 respectively from left to right. Data are presented as Mean ± SEM. (**h**) Representative confocal images showing tubular structures of overexpressed BLOS1 with different tags in mouse primary hepatocytes. (**i**) Tubular BLOS1-GFP expressed in mouse primary hepatocytes distributes on microtubule (indicated by α-Tubulin) (bottom), but not actin filaments (labeled by Phalloidin, top). Merged and single labeling images of magnified insets of boxed areas are shown in bottom panels of each figure. (**j**) co-IP (immunoprecipitation) of BLOS1-HA with co-overexpressed KIF3B-FLAG or KIF13A-FLAG, but not KIF16B-FLAG, in HEK293T whole cell lysate after incubation with anti-FLAG beads, followed by immunoblotting (IB). (**k**) BLOS1 tubules colocalize well with KIF13A-FLAG positive microtubules in mouse primary hepatocytes. Merged and single labeling images of magnified insets of boxed areas are shown on right. Scale bars in all pictures, 10 μm.

Interestingly, when overexpressed in mouse primary hepatocytes, BLOS1 showed a tubular distribution in addition to the scattered puncta (**Fig. 3h**). This observation promoted us to examine the relationship between these tubular structure and cytoskeletons. We found that these tubular structures colocalized well with α-tubulin labeled microtubules (**Fig. 3i**, bottom) but not with actin filaments (labeled with phalloidin) (**Fig. 3i**, top). The microtubule-localized pattern of BLOS1 in primary hepatocytes suggests that BLOS1 may be associated with microtubules.

Since BLOS1 has no reported or predicted microtubule binding motif, we then explored whether other microtubule binding proteins were involved in the microtubular distribution of BLOS1 in primary hepatocytes. Three motor proteins (KIF3B, KIF13A, and KIF16B) from the kinesin superfamily, which have been reported to function in the anterograde transport of vesicles in endocytic system, were taken as candidates. Among these three KIF proteins, KIF3B belongs to the kinesin-2 family, while KIF13A and KIF16B are classified in the kinesin-3 family. The co-immunoprecipitation (co-IP) assays revealed that BLOS1 interacted with both KIF3B and KIF13A, but not KIF16B (**Fig. 3j**). In agreement with these results, we found that the tubular structures that BLOS1 labeled were KIF13A-positive (**Fig. 3k**). Together, these results suggest that BLOS1 may play a role in endosomal trafficking through interacting with KIF proteins on the microtubules.

### Kinesin-3 transports recycling endosome-resident LDLR

We noticed that LDLR mainly localized peripherally underneath the cell membrane in primary hepatocytes (**Fig. 4a**, left), while in Hep G2 cells, LDLR showed a scattered distribution (**Fig. 4a**, right). The differential distribution pattern of LDLR in primary and immortalized cells may reflect the effects of microtubule-localized BLOS1 in primary hepatocytes. Expression of KIF13A-GFP in primary hepatocytes showed the majority of LDLR were enriched at some peripheral domains (**Fig. 4b**), while the KIF13A truncation lacking the motor region (KIF13A-ST) retained LDLR on dispersed vesicles in the cytosol (**Fig. 4c**), suggesting that KIF13A participates in the transport of LDLR to the cell periphery, and dysfunction of KIF13A disturbs the normal trafficking of LDLR.

**Figure 4.**
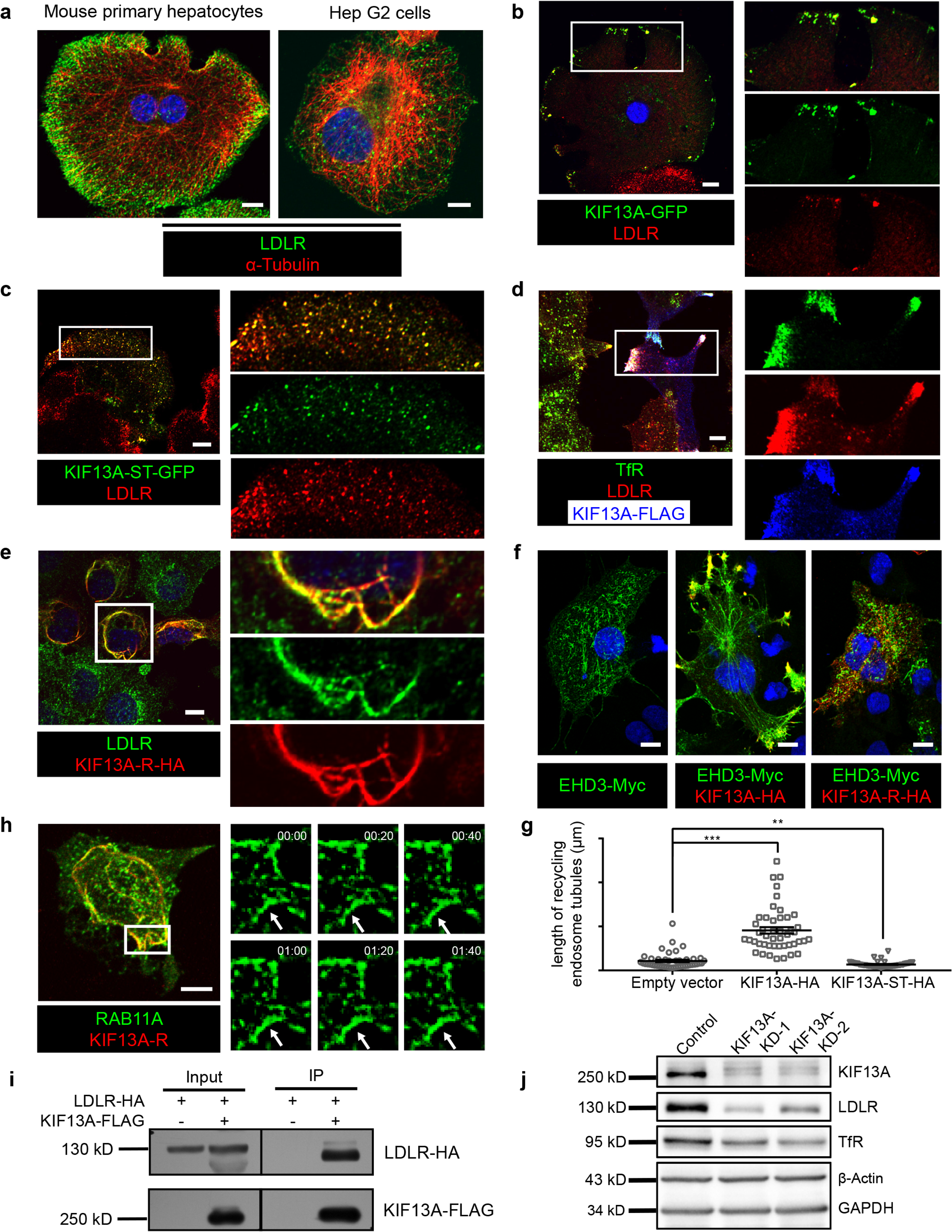
Kinesin-3 transports recycling endosome-resident LDLR. (**a**) Representative confocal images showing different distribution patterns of endogenous LDLR in mouse primary hepatocytes and Hep G2 cells. Microtubules were co-labeled with α-tubulin to show the cell morphology. (**b**) Immunofluorescence staining of LDLR in mouse primary hepatocytes after the expression of KIF13A-GFP. Merged and single labeling images of magnified insets of boxed areas are shown on right. (**c**) Retention of LDLR in the cytosol of mouse primary hepatocytes by expressing KIF13A-ST truncation. Merged and single labeling images of magnified insets of boxed area are shown on right. (**d**) Peripherally distributed LDLR driven by KIF13A-FLAG colocalizes with TfR labeled recycling endosomes in Hep G2 cells. Merged and single labeling images of magnified insets of boxed areas are shown on right. (**e**) Stuck of LDLR on microtubules caused by KIF13A-R point mutant expression in Hep G2 cells. Merged and single labeling images of magnified insets of boxed area are shown on right. (**f**) Representative immunostaining results showing the elongation of recycling endosome tubules caused by full length KIF13A and shortening of recycling endosome tubules caused by KIF13A-ST truncation in Hep G2 cells. (**g**) Average tubular recycling endosomes length measured in empty vector (n=44), KIF13A-HA (n=46) and KIF13A-ST-HA (n=46) expressing cells in (F). Mean ± SEM, two-tailed t test, **p < 0.01, ***p < 0.001. (**h**) Confocal live-cell microscopy of RAB11A-GFP labeled recycling endosomes in KIF13A-R expressing Hep G2 cells. Magnified insets (of boxed area) of consecutive time-lapse images (image/20 s) showed that most of the recycling endosomes were retained on KIF13A-R positive microtubules (see Video 1). White arrows indicate representative recycling endosomes that were almost static during the imaging period. Time stamps are in the format of minutes: seconds. (**i**) Co-IP of LDLR-HA with co-overexpressed KIF13A-FLAG in HEK293T whole cell lysate after incubation with anti-FLAG beads, followed by immunoblotting. (**j**) Immunoblot of LDLR and TfR in KIF13A stable knockdown cells. Scale bars in all pictures, 10 μm. See also **Fig. S3** and **Video 1**.

As BLOS1 is seen on endosomal structures in Hep G2 cells (**Fig. 3**), LDLR was localized to the cell periphery in full-length KIF13A-FLAG overexpressed Hep G2 cells, and these LDLR signals colocalized with the RE marker TfR (**Fig. 4d**), suggesting that KIF13A functions on REs during LDLR recycling. Besides, the rigor mutant of KIF13A (KIF13A-R), an ATPase activity-lacking point mutant which binds to microtubule but is unable to move along the microtubule (Guardia et al., 2016; Nakata & Hirokawa, 1995), anchored LDLR and REs on KIF13A-R positive microtubules (**Fig. 4e**, **Fig. S3a** and **Video 1**). This suggests a critical role of KIF13A in RE-dependent LDLR transport. As a negative control, neither full-length KIF13A nor KIF13A-R mutant affects the cellular distribution of lysosome/MVB marker CD63 (**Fig. S3b,c**).

In agreement with a previous report that KIF13A functions in RE tubule morphogenesis (Delevoye et al., 2014), overexpressed full-length KIF13A elongated tubular REs (labeled by EHD3), while KIF13A-ST truncation had an opposite effect (**Fig. 4f, g**). Live-cell imaging revealed that expression of KIF13A-R affects the movement of REs (labeled by RAB11A), most REs distributed along the KIF13A-R positive microtubules and were almost static during the 2-min imaging period (**Fig. 4h**), indicating that the LDLR anchored on microtubules through KIF13A-R on REs. Furthermore, we found that KIF13A interacted with LDLR in the co-IP assays (**Fig. 4i**), and knockdown of *KIF13A* resembles the reduction of LDLR and TfR occurred in BLOS1 deficiency (**Fig. 4j**). From these results, we firstly report that LDLR in REs is a cargo transported by KIF13A, and dysfunction of KIF13A affects the cellular distribution and homeostasis of LDLR on plasma membrane.

### BLOS1 acts as an adaptor protein of kinesin-2 in the regulation of RE trafficking

We observed that KIF3B exhibited a mostly diffuse distribution when expressed in Hep G2 cells, with no significant effect on the scattered puncta pattern of LDLR (**Fig. 5a**). The expression of KIF3B rigor mutant (KIF3B-R) did not affect the cellular localization of LDLR on REs (labeled by TfR) (**Fig. 5b**). However, by live-cell imaging analysis, we found that, unlike the active movement in control cells, most of the RE puncta in KIF3B-R expressing cells were anchored at dispersed sites on KIF3B-R positive microtubules (**Fig. 5c** and **Video 2**), which suggests that KIF3B may play a role in the trafficking of REs.

**Figure 5.**
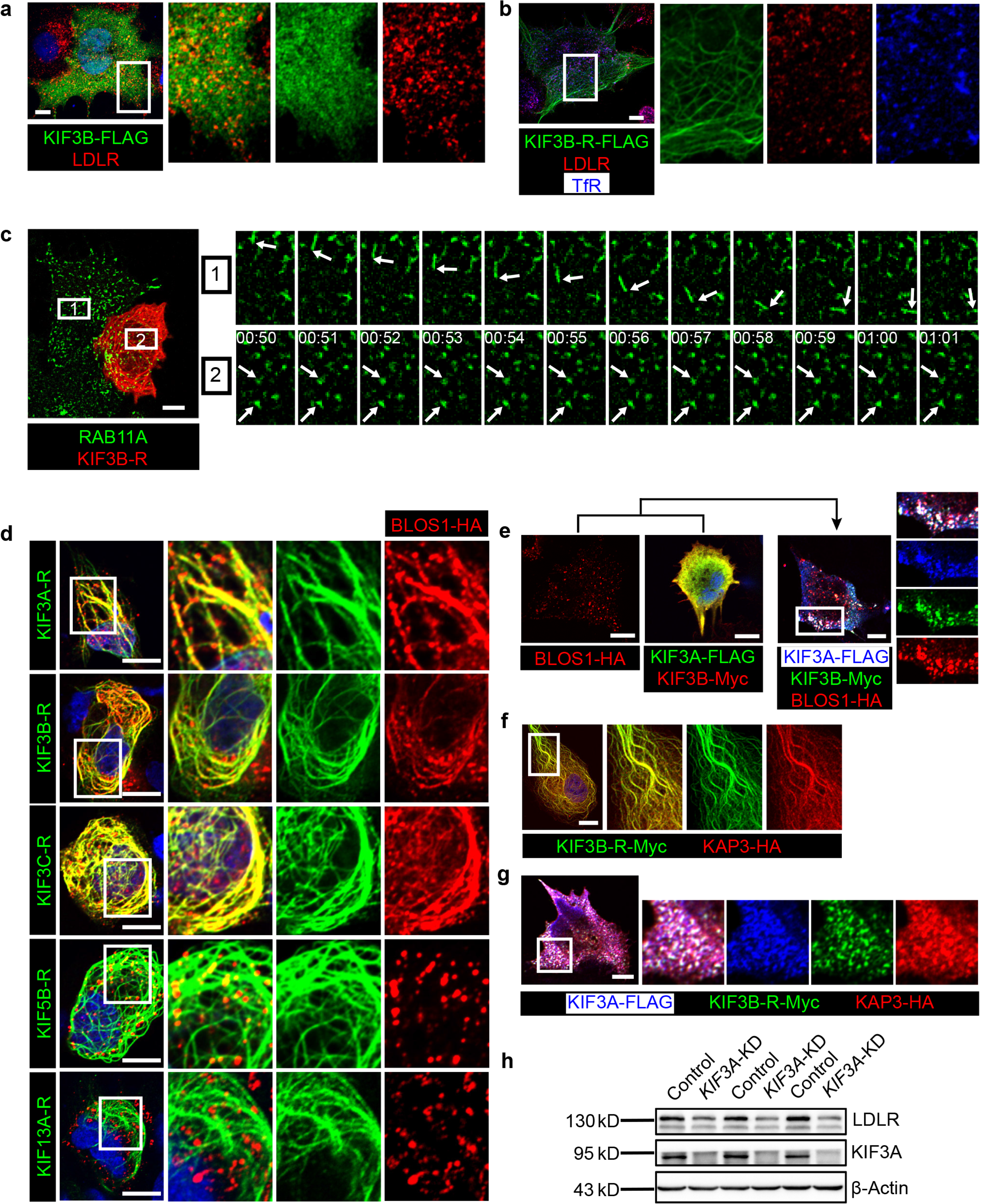
BLOS1 acts as an adaptor protein of kinesin-2 in the regulation of RE anterograde transport. (**a**) Immunofluorescence staining of LDLR in KIF3B-FLAG expressing Hep G2 cells (If not specified, cells used below were all Hep G2 cells). Merged and single labeling images of magnified insets of boxed areas are shown on right. (**b**) Puncta distribution of LDLR and TfR in KIF3B-R expressing cells. Merged and single labeling images of magnified insets of boxed area are shown on right. (**c**) Confocal live-cell microscopy of RAB11A-GFP labeled recycling endosomes in KIF3B-R expressing and non-transfected cells. Magnified insets (of box 1 and 2 in the same time points) of consecutive time-lapse images (image/1 s) showed that recycling endosomes in control cells had active motion, while this movement was impaired in KIF3B-R expressing cells (see Video 2). White arrows indicate representative recycling endosomes in control and KIF3B-R expressing cells. Time stamps are in the format of minutes: seconds. (**d**) Retention of BLOS1 on microtubules caused by co-expressed KIF3A, KIF3B and KIF3C rigor mutants, but not KIF5B or KIF13A rigor mutants. Merged and single labeling images of magnified insets of boxed areas are shown on right. (**e**) Co-expression of BLOS1 with both KIF3A and KIF3B redistributed KIF3A/B to BLOS1 positive puncta. Merged and single labeling images of magnified insets of boxed area are shown on right. (**f, g**) Similar retention of KAP3 on microtubules caused by KIF3B-R (f) and redistribution of KIF3A/B resulting from the co-expression of KAP3 (g). Merged and single labeling images of magnified insets of boxed area are shown on right. (**h**) Immunoblot of LDLR in KIF3A stable knockdown cells (KIF3A-KD cells). Scale bars in all pictures, 10 μm. See also **Video 2**.

In addition, we observed that BLOS1 could be recruited to microtubules by KIF3B-R, and the rigor mutants of two KIF proteins of kinesin-2 (KIF3A or KIF3C) both had the same effect, while KIF13A-R, which also interacted with BLOS1, or KIF5B-R (rigor mutant of a kinesin-1 component) did not affect BLOS1 localization (**Fig. 5d**). Furthermore, when KIF3A and KIF3B, which forms a heterodimer, were co-transfected with BLOS1 in Hep G2 cells, these two KIF proteins were redistributed to BLOS1 positive puncta structures (**Fig. 5e**). One possible explanation for these results is that BLOS1 may form a tight protein complex with KIF3A/B or KIF3A/C heterodimers.

Due to the lack of a cargo-binding region in KIF proteins of kinesin-2, adaptor proteins are required to form a functional kinesin-2 complex for cargo transport. Therefore, we cloned a known kinesin-2 adaptor protein gene *KAP3* and found that when co-expressed with KIF3B-R or KIF3A/B heterodimer, KAP3 show similar localization pattern as BLOS1 (**Fig. 5f, g**), indicating that BLOS1 may act as a new adaptor protein of kinesin-2.

To avoid a possible redundant function between KIF3B and KIF3C, we chose the common subunit KIF3A as the target to be knocked down. Immunoblotting assays revealed that KIF3A knock-down also caused decreased LDLR levels in Hep G2 cells (**Fig. 5h**). Together, our data indicate that BLOS1 may be a new adaptor protein participating in the assembly of functional kinesin-2 complexes, which regulate RE trafficking at dispersed sites on microtubules.

#### Kinesin-2 is essential for the long-range anterograde transport of REs driven by kinesin-3

To explore the detailed mechanism underlying the regulation of RE trafficking by kinesin-2, we labeled REs in both control and *KIF3A*-KD cells with RAB11A-GFP and then analyzed their motion using live-cell imaging. We found that in these cells, the movement of REs could be categorized into two groups, a portion of REs was static or showed slightly local motion, while the other REs moved quickly in anterograde or retrograde transport directions. We named these quickly moving REs as “active REs”. Further analysis revealed that the motion of anterograde transported active REs in *KIF3A*-KD cells was significantly different from that of control cells (**Fig. 6a-d**). In control cells, active REs in anterograde transport sometimes paused and then usually continued their motion in their original anterograde direction (**Fig. 6a** and **Video 3**). In contrast, after the knockdown of *KIF3A* (**Fig. 6b** and **Video 3**) or *BLOC1S1* (**Fig. 6c** and **Video 4**), active REs frequently moved backward after paused anterograde transport. Most of these REs moved back to the perinuclear endocytic recycling compartment (ERC) of their origin by following the backward movement (**Video 3**), suggesting a reduction in the percentage of REs that could eventually be destined to the cell periphery.

**Figure 6.**
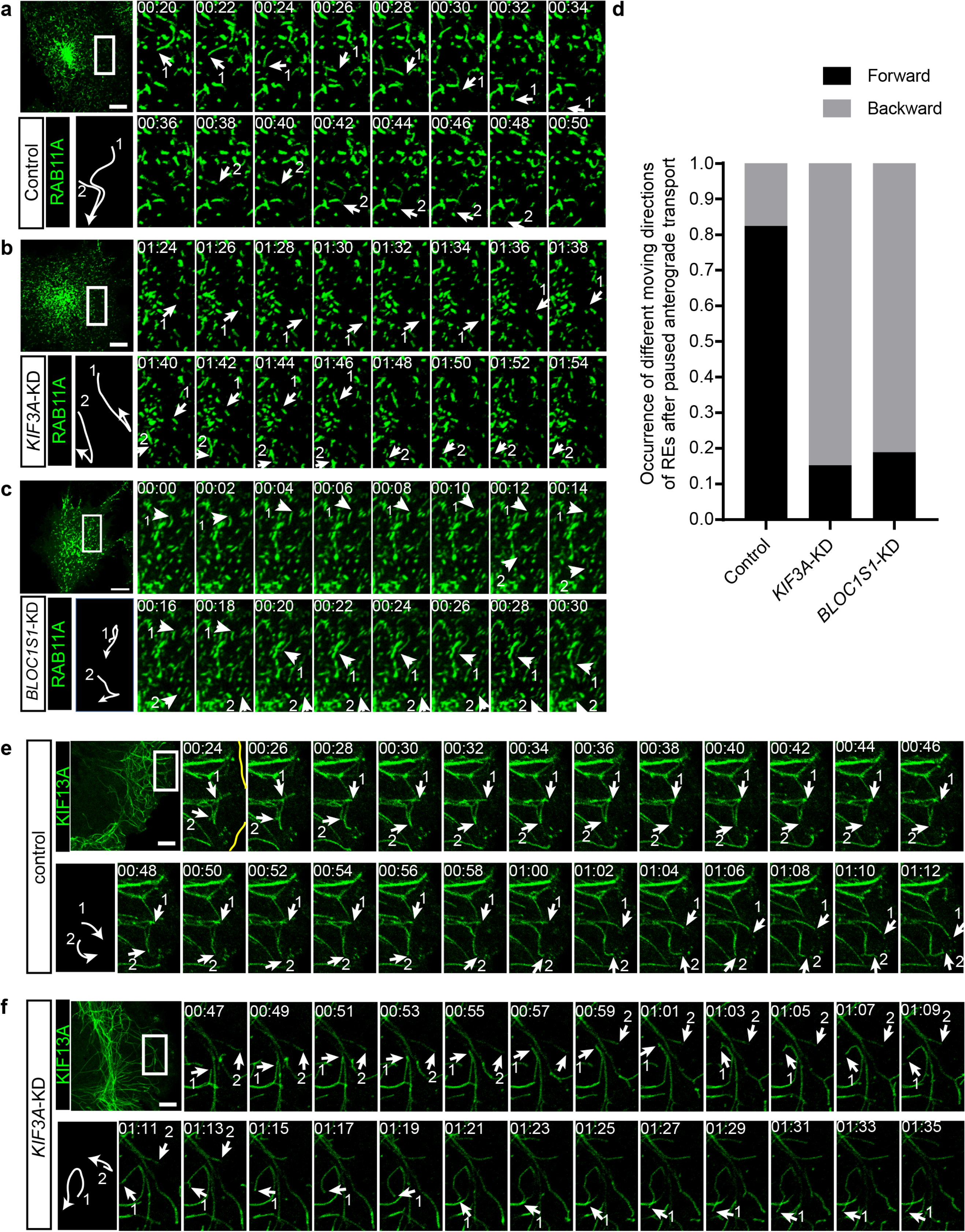
Kinesin-2 is essential for the long-range anterograde transport of REs driven by kinesin-3. (**a-c**) Confocal live-cell microscopy of RAB11A-GFP labeled recycling endosomes in control (a), *KIF3A*-KD (b) and *BLOC1S1*-KD (c) cells (see also Video 3 and Video 4). Magnified insets (of boxed areas) of consecutive time-lapse images (image/2 s) showed that, compared to control cells, the occurrence of backward movement of recycling endosomes following paused anterograde transport is significantly increased in *KIF3A*-KD and *BLOC1S1*-KD cells. White arrows labeled 1 and 2 indicate representative recycling endosomes in these cells. The arrows were drawn perpendicularly to the direction of motion and were reversed when moving in the opposite direction. A trajectory diagram of the representative recycling endosomes is shown on the left bottom. Time stamps are in the format of minutes: seconds. (**d**) Histogram showing the occurrence of forward and backward motion of recycling endosomes following paused anterograde transport in control (n = 50) and *KIF3A*-KD (n = 124) cells. (**e, f**) Confocal live-cell microscopy of KIF13A-GFP tubules in control and *KIF3A*-KD cells (see Video 4). Magnified insets (of boxed areas) of consecutive time-lapse images (image/2 s) showed impaired long-range transport of KIF13A tubules in *KIF3A*-KD cells. White arrows labeled 1 and 2 indicate tips of representative KIF13A-positive tubules in control (e) and *KIF3A*-KD (f) cells. The arrows were drawn perpendicularly to the direction of motion and were reversed when moving in the opposite direction. A trajectory diagram of the representative KIF13A tubule tips is shown on the left bottom. Time stamps are in the format of minutes: seconds. Scale bars in all pictures, 10 μm. See also **Fig. S4**, **Video 3, 4, 5** and **6**.

The above results showed that both kinesin-2 and kinesin-3 participate in the transport of REs. We wondered whether they cooperate in this process. To test whether kinesin-2 is required for the normal function of kinesin-3, we expressed KIF13A-GFP in control and *KIF3A*-KD cells and then analyze the behaviors of these KIF13A-positive tubular structures before and after KIF3A deficiency. We found that, unlike the successful extension to the cell periphery in control cells (**Fig. 6d** and **Video 5**), most of the tips of KIF13A-positive tubules in *KIF3A*-KD cells showed interrupted movement when they encountered other KIF13A tubules (**Fig. 6e** and **Video 5**). Some KIF13A tubule tips moved forward and backward repeatedly on the same track (**Fig. 6f**, arrow 2), while the others paused and then moved backward on another track resembling the behavior of REs in KIF3A-KD cells (**Fig. 6f**, arrow 1). In addition, KIF3A-KD cells showed normal microtubule architecture (**Fig. S4a**) and dynamics (indicated by motility of EB1-GFP positive microtubule tips) (**Fig. S4b** and **Video 6**), suggesting unaffected microtubule network after KIF3A deficiency.

In line with the above results, when KIF13A-R and KIF3B were co-expressed, the distribution of LDLR is determined by KIF13A-R, indicating that KIF13A could function upstream of KIF3B (**Fig. S4c**). However, when KIF3A, KIF3B, and BLOS1 were co-overexpressed to form an intact kinesin-2 complex, peripheral accumulation of LDLR which occurred in the case of KIF13A expression, was not observed (**Fig. S4d**), suggesting that kinesin-2 itself may not drive the long-range anterograde transport of REs.

To further determine the distribution of KIF13A-R and KIF3B-R on microtubules, we co-stained these two rigor mutants with antibodies to acetylated and tyrosinated tubulin, as it has been shown that tyrosinated tubulin has a broader distribution than centrally located acetylated tubulin (Guardia et al., 2016), which resembles the distribution pattern of KIF3B-R and KIF13A. We found that both KIF13A-R and KIF3B-R located on acetylated microtubules (**Fig. S4e, f**, top), and KIF3B-R but not KIF13A-R was additionally found on peripherally distributed tyrosinated microtubules (**Fig. S4e, f**, bottom). These observations indicated that KIF13A and KIF3B may walk along the same set of microtubule tracks with acetylated tubulin and post-translational modifications of tubulin may contributes to the association of KIF13A and KIF3B with microtubules.

### Kinesin-2 and BLOS1 function at specific microtubule intersections

It has been reported that cargoes (lysosomes) pause at intersection points between multiple microtubules and switch to another microtubule track at some specific intersections, and these microtubule intersections serve as important sites for direction determination during cargo transport (Bálint, Verdeny Vilanova, Sandoval Álvarez, & Lakadamyali, 2013; Bergman et al., 2018; Verdeny-Vilanova et al., 2017). We found that, after knockdown of *KIF3A*, when encountering other KIF13A-GFP positive tubules, tips of KIF13A tubules were unable to pass through and move on (**Fig. 7a, b**). As the KIF13A-positive tubules move along microtubules, the site where a tip of KIF13A tubule encounters another KIF13A tubule may represent the intersection point between these two microtubules (**Fig. 7a**). This raised the possibility that the BLOS1-dependent kinesin-2 complex may function at specific microtubule intersections.

**Figure 7.**
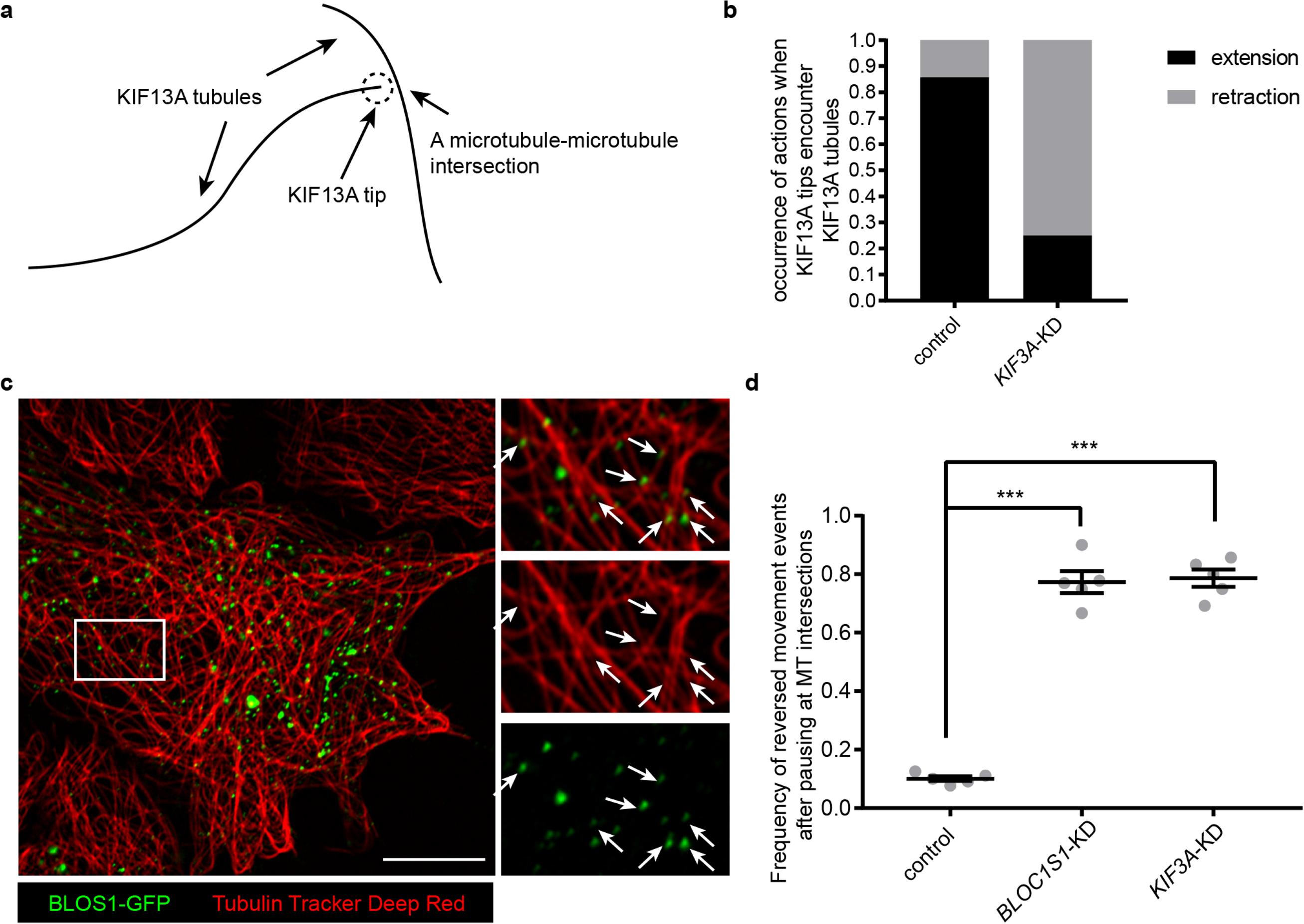
Kinesin-2 and BLOS1 function at specific microtubule intersections. (**a**) Schematic of tip of one KIF13A-GFP tubule encounters another KIF13A-GFP tubule. (**b**) Histogram showing the occurrence of extension and retraction action when tips of KIF13A tubule encounter other KIF13A tubules in control (n = 64) and *KIF3A*-KD (n = 100) cells. (**c**) Super-resolution microscopy of the BLOS1-GFP puncta (green) on microtubule tracks (red, labeled by Tubulin Tracker Deep Red). Merged and single labeling images of magnified insets of boxed area are shown on right. Arrows indicate BLOS1 puncta that locate near microtubule intersections. (**d**) Frequency of reversed movement after pausing at specific microtubule intersection was increased in both *BLOC1S1*-KD and *KIF3A*-KD cells as compared to control cells. Data are presented as Mean ± SEM, n = 5 cells. Two-tailed t test, ***p < 0.001. Scale bars in all pictures, 10 μm. See also Video 7 and 8.

To test this idea, we first detected the distribution of BLOS1 on microtubules using super-resolution microscopy and found that a certain percentage of BLOS1 puncta (77.6 ± 4.6%, n = 8 cells, mean ± SD) located at the intersections of microtubules (**Fig. 7c**), indicating that kinesin-2 may function at these sites. Then we observed the behavior of recycling endosomes (indicated by RAB11A-GFP) on microtubule tracks (stained by Tubulin Tracker Deep Red) in live cells using Zeiss LSM 880 system’s Airyscan super-resolution module. We found that a large proportion of pausing events of recycling endosomes occurred at microtubule intersection sites, and in most cases recycling endosomes moved on without changing their motion polarity after the pausing (**Video 7**). But under certain conditions, recycling endosomes reversed their moving direction after their pausing at specific microtubule intersections (**Video 8**). Furthermore, statistic results showed that the frequency of reversed movement of recycling endosomes after paused anterograde transport at specific microtubule intersections was increased by knockdown of either *BLOC1S1* (77.3 ± 8.4%) or *KIF3A* (78.7 ± 6.6%) as compared to control cells (10.1 ± 1.8%), which means that dysfunction of BLOS1-dependent kinesin-2 complex alters the behavior of recycling endosomes at specific microtubule intersections by changing their motion polarity and finally reduces the recycling of LDLR.

In summary, our observations suggest that BLOS1 acts as an adaptor for kinesin-2 to assist the LDLR cargo transportation driven by kinesin-3 at specific microtubule-microtubule intersections where hurdles may preclude the cargo for smooth long-range anterograde transport. Our results reveal a novel function of BLOS1 in mediating a kinesin switch at microtubule intersections.

## DISCUSSION

Our results support a model in which kinesin-2 (and its adaptor protein BLOS1) functions at checkpoints on microtubules during anterograde transport of recycling endosomes driven by kinesin-3 (**Fig. 8**). The role of BLOS1 in this process is considered to be independent of BLOC-1 or BORC for the following reasons: (1) neither BLOC-1-deficient *pa* mice nor BORC-deficient *Kxd1*-KO mice showed reduced LDLR level; (2) *KIF3A*-KD cells resembled BLOS1 deficient phenotype both in LDLR level and abnormal movement of REs, indicating a kinesin-2-related function. But it remains possible that BLOS1 deficiency caused by dysfunction of BORC may further enhance LDLR degradation due to the proximity of ERC and perinuclearly clustered lysosomes since it has been reported that clustering LEs/lysosomes at the MTOC would generally enhance aggregate degradation and macroautophagy (Bae, Moore, Mella, Hayashi, & Hollien, 2019; Korolchuk et al., 2011).

**Figure 8.**
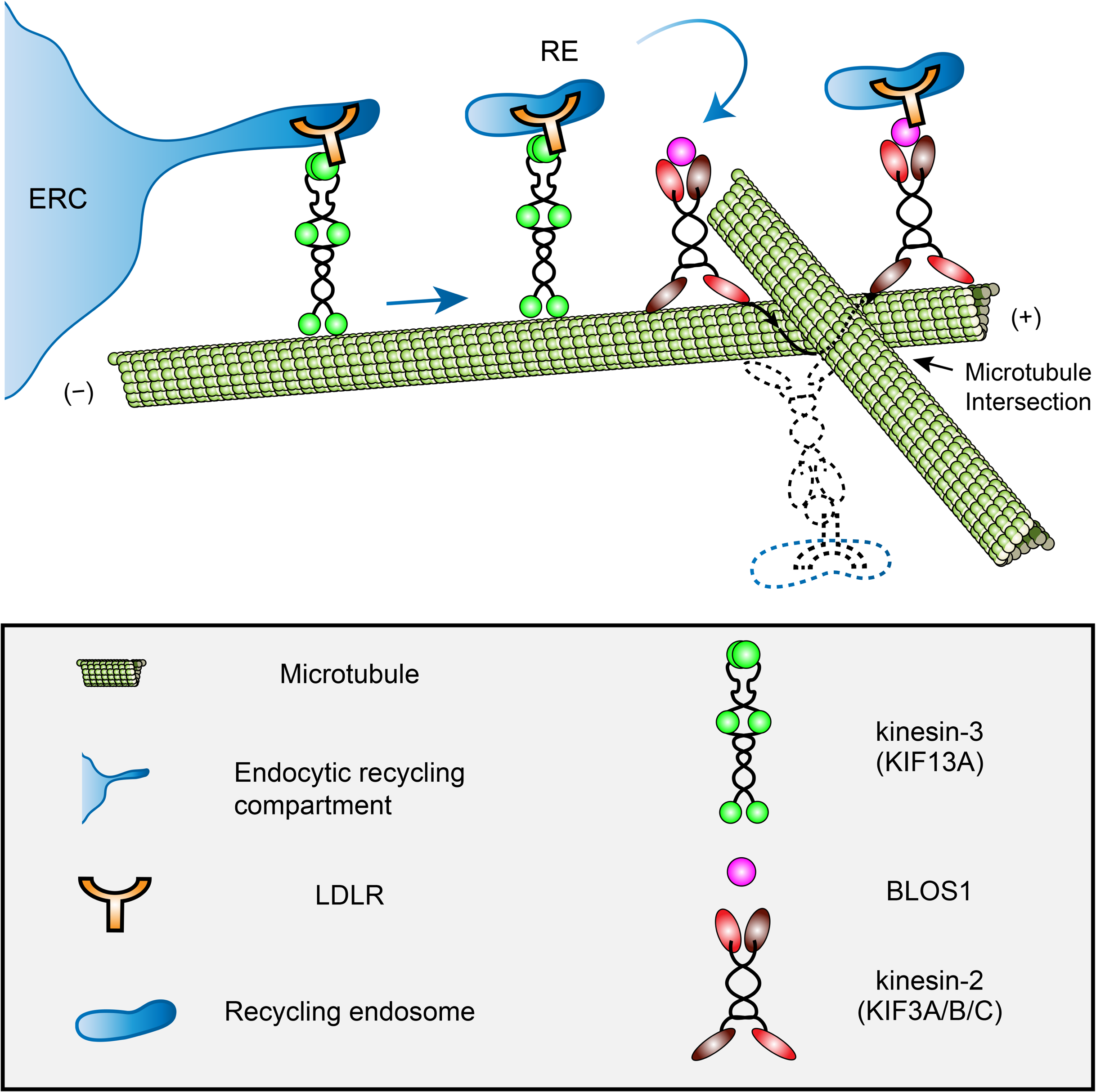
A proposed model of how BLOS1 coordinates kinesin-3 and kinesin-2 at microtubule intersections. KIF13A drives the generation and extension of tubules at the initial peripheral transport of recycling endosomes (REs) from the endocytic recycling compartment (ERC). When REs (and associated LDLR) arrive at specific microtubule-microtubule intersections, their motion will be paused by steric hindrance. At these sites, through the interaction with kinesin-3 (KIF13A) and LDLR, BLOS1 (and kinesin-2 heterodimers) assists the REs in overcoming obstacles probably by switching between microtubule protofilaments and finally causing the rotation of cargo along the microtubule surface. Dysfunction of kinesin-2 results in the inability of REs to pass through these sites and frequently backward movements at specific microtubule intersections. Thus, impaired recycling causes alternative lysosomal degradation of LDLR. See also Video 9.

As for the complexity in kinesin motors and cargoes, it has been reported that one kinesin could use different adaptor proteins for various cargoes and different kinesins may transport one cargo on distinct microtubule tracks (Guardia et al., 2016; Hirokawa et al., 2009). We found here that kinesin-2 functions downstream of kinesin-3 in the same trafficking pathway of REs. This observation expands knowledge of how cargoes are transported by kinesins.

Previous *in vivo* and *in vitro* studies have shown that microtubule intersections may serve as the pausing site for cargoes transport by kinesins when the size of cargoes are comparable to intersection spacing (Bálint et al., 2013; Verdeny-Vilanova et al., 2017), and it has been suggested that both the intersection geometry and motor number influence cargo routing at these intersections (Bergman et al., 2018; Erickson, Gross, & Yu, 2013). In addition, members of the heterodimeric kinesin-2 subfamily switch protofilaments during their processive movements, which may contribute to the overcoming of microtubule obstacles (Brunnbauer et al., 2012; Hoeprich, Mickolajczyk, Nelson, Hancock, & Berger, 2017). Such off-axis mode motion of cargoes were also observed at microtubule intersections *in vivo* (Verdeny-Vilanova et al., 2017). Therefore, in the context of recycling endosome trafficking, kinesin-2 may be used to pass through these intersections by protofilament switching. In agreement with this hypothesis, we observed that BLOS1 puncta showed similar off-axis mode motion during the time period of live-cell imaging (**Video 9**). Although we found that kinesin-2 (and BLOS1) is essential for the circumvention of these intersections after paused anterograde transport of recycling endosomes, the detailed mechanism needs further investigation. It may involve other microtubule-associated proteins and the geometry change of microtubule intersections.

We recently reported the lipidomic profiling of *pa* mice (another BLOC-1 subunit mutant) (Ma et al., 2019) by showing an anti-anthrogenic effect. Regarding the dyslipidemic profile in BLOS1-hepatocyte specific knockout mice, its clinical relevance may not be exactly mimicking patients with recessive *BLOC1S1* mutations as BLOS1 deficiency is an embryonic lethal (Scott et al., 2014; Zhang et al., 2014), and the whole body knockout of BLOS1 could be different given some hypomorphic mutations are viable. Nevertheless, the involvement of BLOS1 in LDLR endosomal recycling may provide insights into the general regulatory mechanism of LDLR trafficking and the understanding of impaired LDL clearance in the liver.

## MATERIALS AND METHODS

### Key resources

All resources used in this study are listed in **Table S1**.

### Experimental model and subject details

#### Animals

The *Bloc1s1* floxed mice (loxp mice) and *Kxd1*-KO mice were originally generated in our lab(Yang et al., 2012; Zhang et al., 2014) and subsequently bred locally in the animal facility (specified pathogen free) of Institute of Genetics and Developmental Biology, Chinese Academy of Sciences. *pa* mice were derived from The Jackson Laboratory. Alb-Cre tool mice were supplied by Model Animal Research Center of Nanjing University. To generate liver-specific *Bloc1s1* knockout mice (cKO mice), Alb-Cre mice were crossed with loxp mice. Colonies were maintained by breeding Alb-Cre; loxp/loxp mice with loxp/loxp mice. Control loxp mice and cKO mice were littermates. Male mice of 3 months were used unless is stated. Mice were kept under a 12 h dark-light period and provided a standard chow diet. For starvation treatment, mice were fasted for 12 h overnight, and all mice livers used for Oil Red O staining were obtained at the next morning. All animal work was approved by the Ethical Review Committee in the Institute of Genetics and Developmental Biology, Chinese Academy of Sciences.

#### Isolation and culture of mouse primary hepatocytes

A two-step collagenase perfusion technique was used to isolate primary hepatocytes from loxp and cKO mice liver as described (Li, Ralphs, & Tosh, 2010). Mice were euthanized by CO_2_ inhalation, the abdominal cavity was opened, and then the precaval vein was closed with a vascular clamp. Liver perfusion from the portal vein was initiated with 50 mL pre-warmed (37℃) HBSS (pH 7.4, no calcium and magnesium) containing 10 mM HEPES and 200 µM EDTA for 10 min. After the liver turned pale, change perfusion medium and perfuse with 50 mL pre-warmed 50 HBSS (pH 7.4, with calcium and magnesium) containing 0.5 mg/mL collagenase IV and 20mM HEPES for 10-15 min. Then remove the entire liver to a petri dish containing HBSS (with calcium and magnesium) on ice, and dissociate the liver lobs by tearing with forceps. The resulting cell suspension was filtered through a 70 µm mesh filter and washed three times with pre-cooled HBSS (centrifuged at 50 × *g* for 2 min). Cells were then suspended in 10 mL William’s E Medium (Gibco) supplemented with 10% (v/v) fetal bovine serum (FBS, Gibco), 1% (v/v) 200 mM GlutaMAX (Gibco) and 1% (v/v) Nonessential amino acids (NEAA, Gibco), and then seeded at a final density of 0.4 × 10^6^ cells per mL onto collagen I-coated dishes (coverslips). Cells were incubated with 5% CO_2_ at 37℃ for 2 h and then remove the medium containing attached cells and replaced with fresh culture medium. Transfections were done 4 h after attachment using jetPEI-Hepatocyte reagent (Polyplus).

#### Culture of cell lines

Hep G2 cells were obtained from the cell bank of the Chinese Academy of Sciences (Shanghai, China). All cell lines used in experiments were maintained at 37℃ with 5% CO_2_. Hep G2 cells were cultured in Minimal Essential Medium (MEM, Hyclone) supplemented with 10% (v/v) FBS, 1% 100 mM sodium pyruvate (Gibco) and 1% NEAA (Gibco).

#### E. coli strain used for GST-fusion protein purification

The strain used for protein expression and purification was *Escherichia coli* (*E. coli*) BL21 (DE3) (Vazyme), and the expressing vector was PGEX-4T-1 (Pharmacia). First, BL21 competent cells were transformed with the expression plasmid and selected on LB plates supplemented with 100 µg/mL ampicillin overnight at 37℃. Single colonies were picked and grown overnight in 5 mL LB medium supplemented with 100 µg/mL ampicillin at 37℃ with shaking at 300 rpm. On next morning, the overnight culture was diluted 1:100 into fresh 100 mL LB medium and cultured at 37℃ (about 2.5 h) until the optical density at 600 nm (OD_600_) reached 0.5 to 0.7. Isopropyl-1-thio-β-D-galactopyranoside (IPTG) was added to final concentration of 0.4 mM and fusion protein expression induced at 25℃ for 8 h.

### Methods

#### Constructs

For the expression of N-terminal tagged FLAG/Myc-fusion proteins (BLOS1-FLAG, BLOS1-Myc), pCMV-Tag-2B/3B vectors (Agilent) were used with cDNA fragments inserted into multiple cloning sites. pEGFP-C2 (Clontech) was used to generate BLOS1-GFP-C2 and RAB11A-GFP plasmids. pEGFP-N2 (Clontech) was used for KIF13A-GFP, KIF13A-ST-GFP expression. For the red fluorescent protein Scarlet(Bindels et al., 2016) expression, ORF of GFP was substituted by the coding sequence of Scarlet by modifying pEGFP-C2 and pEGFP-N2 vectors without changing other DNA elements. Similarly, GFP ORF in the pEGFP-N2 vector was replaced by FLAG/Myc/HA coding sequences to generate plasmid for C-terminal fusion proteins (BLOS1-HA, LDLR-HA, KIF13A-FLAG and all other KIF proteins with corresponding FLAG/Myc/HA tags) expression. The expressing plasmids of KIF protein rigor mutants were generated by site-directed mutagenesis using primers shown in **Table S2**.

The shRNA expressing plasmids used in stable knockdown cell line (*BLOC1S1*-KD, *KIF13A-*KD, and *KIF3A*-KD cells) construction were modified from pSilencer 5.1-H1 Retro vector (Ambion) by introducing a *Mfe*Ⅰ cleavage site (same overhang with EcoRⅠ) between 2166 to 2172 just after the *Xho*Ⅰ site and upstream of the expressing element of shRNA. Take KIF3A knockdown plasmid as an example, three separate shRNA expressing plasmids were generated by annealing the templates (Table S2) and inserting into the modified plasmids. One plasmid was digested with *Xho*Ⅰ and *EcoR*Ⅰ to get the whole shRNA expressing element, another was digested with *Xho*Ⅰ and *Mfe*Ⅰ. Then ligation was performed to generate a new plasmid containing two sets of shRNA expressing elements. This procedure was repeated until all templates for a specific gene were introduced on one plasmid.

ORF of KIF5B was amplified from the cDNA of Hep G2 cells, and all the other ORFs were amplified from the cDNA obtained by reverse transcription of total mRNA of mouse liver. For the construction of KIF3A-FLAG-KIF3B-Myc-BLOS1-HA co-overexpression plasmid, the expression cassette of KIF3B-Myc was amplified and then introduced into *BspT*Ⅰ digested KIF3A-FLAG plasmid by recombination (ClonExpress Ⅱ One Step Cloning Kit, Vazyme) after which the *BspT*Ⅰ site was kept intact, and the expression cassette of BLOS1-HA was inserted by a similar procedure.

#### Immunoblotting

Homogenates of either cell cultures or tissues were used in western blot analysis. Cells in one well of 6 well plates were collected in ice-cold PBS (Gibco) and lysed in 200 µL lysis buffer (50 mM Tris-HCl pH 7.0, 150 mM NaCl, 1 mM EDTA, 1% Triton X-100) supplemented with protease inhibitor cocktail. Tissue samples (about 50 mg) were homogenized with a micro tissue grinder in 500 µL lysis buffer. All homogenates were rotated for 1 h at 4℃ in a vertical rotator and centrifuged at 13000 × *g* for 15 min. Supernatants with soluble proteins were mixed with 6 × Laemmli loading buffer with β-mercaptoethanol and boiled for 5 min. Protein samples were resolved by SDS-PAGE on 10% or 15% Tris-Glycine buffered polyacrylamide gels and transferred to PVDF membranes using mini trans-blot module (Bio-Rad).

The blots were blocked in 5% (w/v) non-fat milk (BD) at room-temperature for 1 h in PBS/0.1% (v/v) Tween 20 (PBST). After a brief wash in PBST, membranes were then incubated in primary antibody diluted in 3% (w/v) bovine serum albumin (BSA) dissolved in PBST overnight at 4℃. On the following day, the membranes was washed three times, each time for 10 min in PBST with shaking and then incubated with 1:5000 HRP-conjugated secondary antibodies (Zsbio) diluted in blocking buffer at room temperature for 1 h. Three more 10 min washes with PBST were then performed before detection using Chemiluminescent Substrate (Thermo) and imaging with a Minchemi system (Sage Creation). Quantification was performed using Fiji(Schindelin et al., 2012). The following primary antibodies were used (additional information was shown in the Table S1): anti-LDLR (1/5000 for ab52818; 1/2000 for ab30532), anti-TfR (1/2000), anti-β-Actin (1/50000), anti-KIF13A (1/1000), anti-KIF3A (1/1000), anti-Pallidin (1/2000), anti-Dysbindin (1/20000), anti-GST (1/5000), anti-FLAG (Sigma, Cat#F3165, 1/5000), anti-Myc (MBL, Cat#562, 1/2000), anti-HA (Abcam, 1/5000).

#### Co-immunoprecipitation

All immunoprecipitations were performed in the absence of cross-linking reagents. Cells in 6 well plates were transfected with FLAG empty vector/FLAG fusion protein plasmid and corresponding candidates of interacting protein plasmids for 48 to 72 h, and cell lysates were prepared in cell lysis buffer as described above. 40 µL anti-FLAG M2 agarose beads (Sigma) were used for the enrichment of FLAG-fusion proteins of each well by incubating with the supernatants overnight at 4℃ on a vertical rotator. Sufficient washes were carried out at 4℃ using lysis buffer with a total time of 30 min on a vertical rotator (5 to 6 washes), beads were pelleted by centrifuging at 5000 × *g* for 30 s after each wash. Proteins were eluted by boiling with 50 µL 2 × Laemmli loading buffer for 5 min and subsequent western blot was performed as described above. For the detection of eluted FLAG-fusion proteins, a particular HRP-conjugated secondary antibody (Abcam, Cat# ab131368) which preferentially detects the non-reduced form of mouse IgG over the reduced, SDS-denatured forms was used to eliminate the IgG band results from the anti-FLAG M2 beads. ***GST pull-down assay*** The full length and truncated BLOS1s were expressed in *E. coli* BL21 as described above, and GST protein which was induced at 37℃ for 4 h served as control. After induction, bacteria were collected by centrifuge at 4000 × *g*, 4℃ for 15 min. The bacteria pellet was resuspended in 10 mL ice-cold lysis buffer (50 mM Tris-HCl pH 8.0, 150 mM NaCl, 5 mM EDTA, 1% Triton X-100) supplemented with protease inhibitor and 1 mM DTT. The suspension was then sonicated on ice using a probe-tip sonicator using the following parameters: 10 s each with 30 s rest at 200 W for a total time of 15 min. The supernatant was collected and stored at -20℃ after centrifugation at 12,000 × *g*, 4℃ for 15 min.

For GST pull-down assay, 50 µL GST protein-containing supernatant diluted in 450 µL lysis buffer was used to incubate with 50 µL slurry of glutathione-Sepharose beads at 4℃ for 6 h, and the dosage of supernatants of other GST fusion proteins was determined by Coomassie brilliant blue staining. The beads were pelleted at 5000 × *g* for 30 s and washed 5 to 6 times with lysis buffer before the incubation with 500 µL mouse liver lysate overnight at 4℃. After incubation, beads were washed with tissue lysis buffer for another 5 to 6 times, and binding proteins were eluted with 2 × Laemmli loading buffer by boiling for 5 min. Immunoblots were performed as described above.

#### Oil Red O staining of mouse liver sections

Freshly collected liver lobes of control (loxp) and cKO mice fed in chow diets or under starvation were embedded in OCT (Tissue-Tek) and flash-freezed immediately with liquid nitrogen. Samples were left in the chamber of the freezing microtome with the temperature set at -20℃ for at least 1 h before the sectioning. Three consecutive 10-µm-thick sections of the liver were collected onto one slide and fixed immediately in ice-cold 4% (w/v) paraformaldehyde dissolved in PBS (pH 7.4) for 10 min and then rinsed in three changes of distilled water. Slides were dehydrated in 60% isopropyl alcohol for 2 min before the immersed by Oil Red O staining solution (1.5 part of Oil Red O saturated isopropyl alcohol solution mixed with one part of distilled water) at room temperature for 15 min. Slides were rinsed in 60% isopropyl alcohol for two times and then placed in distilled water before the next step. Counterstaining of the nucleus with Mayer′s hematoxylin was done by submerging the sections in hematoxylin for 10 s and thereafter dipping the sections in distilled water for three times before bluing the stain in PBS (pH 7.2) for 5 min. Slides were mounted in water-soluble mounting medium (PBS: glycerol =1:9), and coverslip edges were sealed with nail polish. Micrographs were acquired using an Olympus DP71 imaging system, and Fiji was used in the quantification of Oil Red O staining areas.

#### Immunocytochemistry and immunofluorescence imaging

Coverslips were pre-coated with collagen Ⅰ (Sigma) for Hep G2 attachment. Cells attached on coverslips were fixed in 4% (w/v) formaldehyde dissolved in PBS and immunocytochemistry (ICC) was performed according to a general ICC protocol (Abcam). Permeabilization of cells was done by incubating in PBS containing 0.1% (v/v) Triton X-100 for 15 min. Blocking was done with 3% (w/v) BSA in PBS. The following primary antibodies were used (additional information was shown in Table S1): anti-mouse LDLR (R&D, Cat#AF2255, 1/100), anti-human LDLR (R&D, Cat#AF2148, 1/100), anti-TfR (1/200), anti-EEA1 (1/500), anti-Cytochrome C (1/250), anti-CD63 (1/200), anti-FLAG (Sigma, Cat#F3165, 1/1000; Sigma, Cat#F7425, 1/1000), anti-Myc (MBL, Cat#562, 1/500; MBL, Cat#M192-3S, 1/500), anti-HA (Santa Cruz, Cat#sc-7392, 1/100; Abcam, Cat#ab9134, 1/1000) and anti-α-Tubulin (Abcam, Cat#ab7291, 1/500; Abcam, Cat#ab18251, 1/1000). Coverslips were incubated with primary antibodies diluted in blocking buffer overnight at 4℃ and wash buffer used after antibody incubation was PBS containing 0.1% (v/v) Tween 20. All Alexa Fluor-conjugated secondary antibodies (Invitrogen/Abcam) were diluted 1/1000 and used with an incubating time of 1 h at 37℃ in the dark. After the wash, coverslips were mounted on glass slides with Prolong Gold Antifade Mountant (Invitrogen), sealed with nail polish and stored in the dark at 4℃ before imaging. Confocal imaging was carried out using a Nikon Eclipse Ti Confocal Laser Microscope System with NIS-Elements Software (Nikon, Japan). Super-resolution images were acquired in a Zeiss LSM 880 system with Airyscan module. Images were analyzed and quantified were Fiji software with raw data imported through Bio-Formats Importer.

#### Electrophoretic separation of plasma lipoproteins in native gradient polyacrylamide gel

Lipoproteins in plasma samples were pre-stained with Sudan Black B (SBB) staining solution (SBB saturated ethanol solution) by mixing 2 µL of SBB staining solution with 30 µL plasma and incubating at 37℃ for 30 min. A 4% to 15% polyacrylamide gradient mini gel prepared with Tris-HCl buffer (pH 8.3) was used to separate lipoproteins, and the running buffer (0.6 g/L Tris, 2.88 g/L Glycine, pH 8.3) was changed several times during the overnight electrophoresis at 4℃ with a voltage of 70 V. For the Oil Red O staining, unstained plasma samples were separated by electrophoresis and then stained with 0.2% (w/v) Oil Red O methanol solution for 30 min with shaking.

#### Isolation of plasma LDL and LDL-DiI endocytosis assay

LDL of pooled mouse plasma were isolated by sequential ultracentrifugation as described(Havel, Eder, & Bragdon, 1955). High-density salt solution (1.346 g/mL) was prepared by dissolving 153 g NaCl and 354 g KBr in Milli-Q water to a total volume of 1L. Low-density salt solution (1.005 g/mL) contained 0.15 M NaCl. Salt solutions of other densities were prepared by mixing the high-density and low-density salt solutions at different ratios. 5 mL pooled plasma with the density of 1.006 g/mL was mixed with 1 mL of 1.085 g/mL salt solution to reach a total density of 1.019 g/mL and ultracentrifuged at 180,000 × *g*, 4℃ for 12 h. The top 1.5 mL layer containing lipoproteins with a density less than 1.019 g/mL (mainly VLDL) was collected, and additional 1.5 mL salt solution (1.200 g/mL) was mixed with the remaining plasma to reach a density of 1.063 g/mL. Ultracentrifugation was performed as above, and the top 1.5 mL solution containing lipoproteins with the density between 1.019 g/mL to 1.063 g/mL (LDL) was collected. All collected lipoproteins were dialyzed in 500 mL 0.15 M NaCl twice at 4℃ before electrophoresis or DiI labeling.

For DiI labeling, 1 mg of purified LDL were mixed with 50 µL DiI stock solution (3 mg/mL in DMSO; Sigma) and incubated at 37℃ for 8 h. The density was adjusted to 1.063 g/mL, and ultracentrifugation was performed. Collected LDL-DiI on the top layer was dialyzed with 0.15 M NaCl and further filter-sterilized (0.1 µm, Millipore). LDL-DiI were added to the medium of primary hepatocytes at the final concentration of 10 µg/mL, and binding of LDL-DiI to the cell surface was done at 4℃ for 30 min. After the binding, cells at different time points post endocytosis were fixed and visualized.

#### Generation of stable knockdown cell lines

Hep G2 cells in 24 well plates were transfected with targeted gene shRNA or scrambled negative control shRNA expressing plasmids and split into 60 mm dishes at the concentration of 10% confluent 24 h post-transfection. The selection was started the day after splitting using 2 µg/mL puromycin (concentration determined by preliminary experiment). Culture medium was changed every day in the first week and every 3 days after that. Cells were grown for 3 weeks under selection pressure, and single colonies were picked and expanded in 24 well plates with puromycin added. Western blot of target proteins was performed to identify positive colonies.

#### Inhibition of lysosomal degradation

Hep G2 cells were plated in 12 well plates at a seeding density of ∼50% confluency and were cultured for 12 h before leupeptin treatment. Leupeptin (Sigma) dissolved in Milli-Q water was added to a final concentration of 50 µM in the culture medium. Cells were collected 24 h later, and western blot was performed with indicated antibodies as previously described.

#### Live cell Imaging and super-resolution microscopy

Hep G2 cells were cultured on collagen I-coated 35 mm µ-Dish (*ibid*) with a high-performance glass bottom (170 ± 5 µm). Hep G2 cells were transfected with GFP or/and Scarlet fusion protein expressing plasmids using lipofectamine 3000 reagent 24 h before the imaging. Live cell imaging was carried out on a Nikon Eclipse Ti Confocal Laser Microscope System equipped with 405 nm (20 mW), 488 (50 mW) and 561 (100 mW) laser lines, temperature controller, a 100 × oil-immersion objective (Nikon, NA = 1.40) and appropriate filter sets. Before imaging, the culture medium was replaced by CO_2_-independent Live Cell Imaging Solution (Invitrogen) and adapted for 15 min. Image series were captured using the 488 laser line (∼10% laser power) and 561 laser line (∼5% laser power) with 1 second interval and total imaging time of 120 s under the “live” mode.

Super-resolution microscopy of microtubules was carried out on Zeiss LSM 880 system equipped with Airyscan module and cell incubation chamber. Cells transfected with RAB11A-GFP were first stained with Tubulin Tracker Deep Red according to the manufacturer’s instruction, and image series were then captured using the 488 laser line and 633 laser line using Airyscan module with a interval time of 0.32 second and zoom factor of 3. Subsequent imaging processing was accomplished using ZEN 2.3 (Zeiss) and Fiji software.

#### Quantification and statistical analysis

Data were presented as Mean ± SEM unless stated, the statistical significance of mean differences was determined using two-tailed Student’s t test as indicated in the figure legends. The sample size (n) was also indicated in the corresponding figure legends, which represents the number of identically-treated replicates. Statistical significance is defined as, n.s., not significant, *p < 0.05, **p < 0.01, ***p < 0.001. Statistical analyses were performed using GraphPad Prism.

## Acknowledgments

We thank Prof. Richard T. Swank for his proofreading this manuscript and comments. This work was partially supported by grants from the Ministry of Science and Technology of China (2019YFA080055), and the National Natural Science Foundation of China (31830054; 91954000; 91539204; 81670789).

## Author contributions

C. Z. and W. L. designed the experiments and wrote the manuscript. C. H. and G. S. analyzed the data and supervised the project. C. Z. performed all the assays.

## Data availability

The authors declare that data supporting the findings of this study are available within the paper and its supplementary information files.

## Supplemental Information

### Brief introduction to supplementary figures & movies

**Fig. S1** (related to **Fig. 1**) shows generation and genotyping of cKO mice, as well as the increased plasma ApoE content in cKO mice identified by mass spectrometry. **Fig. S2** (related to **Fig. 2**) shows the elevated plasma ApoB level in cKO mice and the interaction details between BLOS1 and LDLR detected by co-IP and GST pull-down assays. **Fig. S3** (related to **Fig. 4**) shows that KIF13A and its rigor mutant do not affect lysosome distribution, which is obviously different from the role KIF13A plays on recycling endosomes. **Fig. S4** (related to **Fig. 6**) shows the unaffected microtubule architecture and dynamic in *KIF3A*-KD cells, and both kinesin-2 and kinesin-3 bind to the same subset of microtubule tracks with slightly different.

**Video 1** (related to **Fig. 4**) shows by time-lapse confocal imaging the abnormal tubular distribution and reduced motility of recycling endosomes in KIF13A-R expressing cells. **Video 2** (related to **Fig. 5**) shows by time-lapse confocal imaging the normal puncta distribution and reduced motility of recycling endosomes in cells with KIF3B-R expression. **Video 3** (related to **Fig. 6**) shows by time-lapse confocal imaging the significantly increased the occurrence of backward movement of recycling endosomes following paused anterograde transport in *KIF3A*-KD cells comparing to control cells. **Video 4** (related to **Fig. 6**) shows by time-lapse confocal imaging the similar phenomenon in *BLOC1S1*-KD cells observed in KIF3A-KD cells showed in **Video 3**. **Video 5** (related to **Fig. 6**) shows by time-lapse confocal imaging that, compared to control cells, the KIF13A-positive tubules in *KIF3A*-KD cells were unable to reach the cell periphery with interrupted movement at specific points in their routes. **Video 6** (related to **Fig. 6**) shows by time-lapse confocal imaging that control cells and *KIF3A*-KD cells have similar microtubule growth rate, indicating the unaltered microtubule dynamic after *KIF3A* depletion. **Video 7** (related to **Fig. 7**) shows by time-lapse super-resolution imaging that certain recycling endosomes (marked with magenta circles) paused at specific microtubule-microtubule intersections (indicated by white arrows) and then moved on microtubule tracks without motion polarity changing. **Video 8** (related to **Fig. 7**) shows by time-lapse super-resolution imaging that certain recycling endosomes (marked with magenta circles) paused at specific microtubule-microtubule intersections (indicated by white arrows) and then reversed their motion polarity. **Video 9** (related to **Fig. 8**) shows by time-lapse confocal imaging the off-axis mode motion of BLOS1 puncta during recording period, indicating the protofilament switching along microtubules.

---

**Figure S1.**
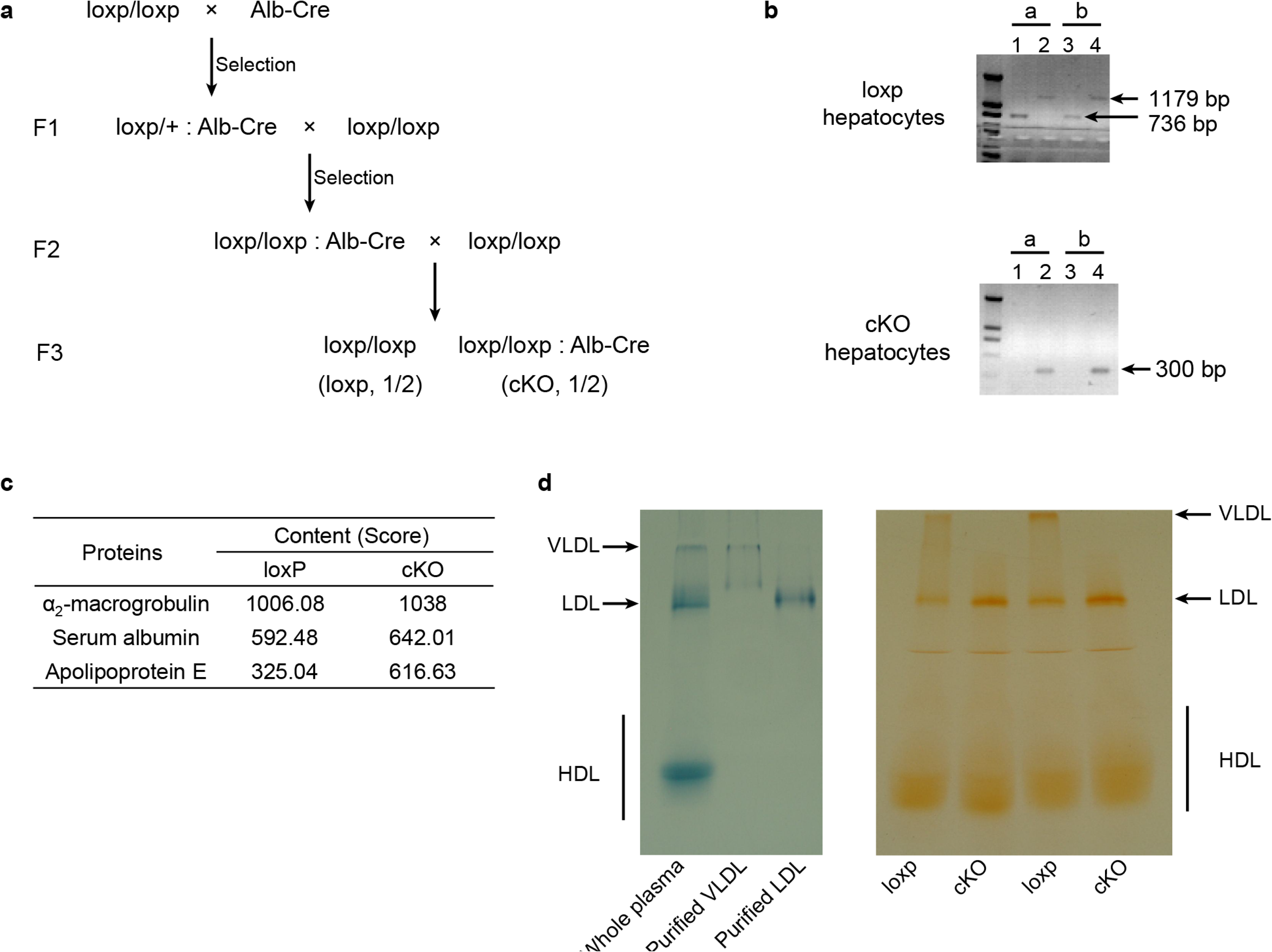
Generation of cKO mice and the abnormal plasma lipoprotein level in cKO mice, Related to**Fig. 1**. (**a**) Breeding strategy for the generation and maintaining of loxp and cKO mice littermates which were used in subsequent experiments. (**b**) Genotyping of hepatocytes of loxp and cKO mice. a and b represent two different mice. Primers used in lane 1 and 3 recognize the loxp sites, and primers used in lane 2 and 4 amplify a larger fragment containing both loxp sites. (**c**) Mass spectrometry of p34 gel slices showing increased ApoE content (indicated by Score) in cKO mice plasma. (**d**) Bands of purified lipoproteins after native gel electrophoresis show the position of these lipoproteins in separated plasma samples (left). Oil Red O staining result of native gel separated unstained lipoproteins (right).

**Figure S2.**
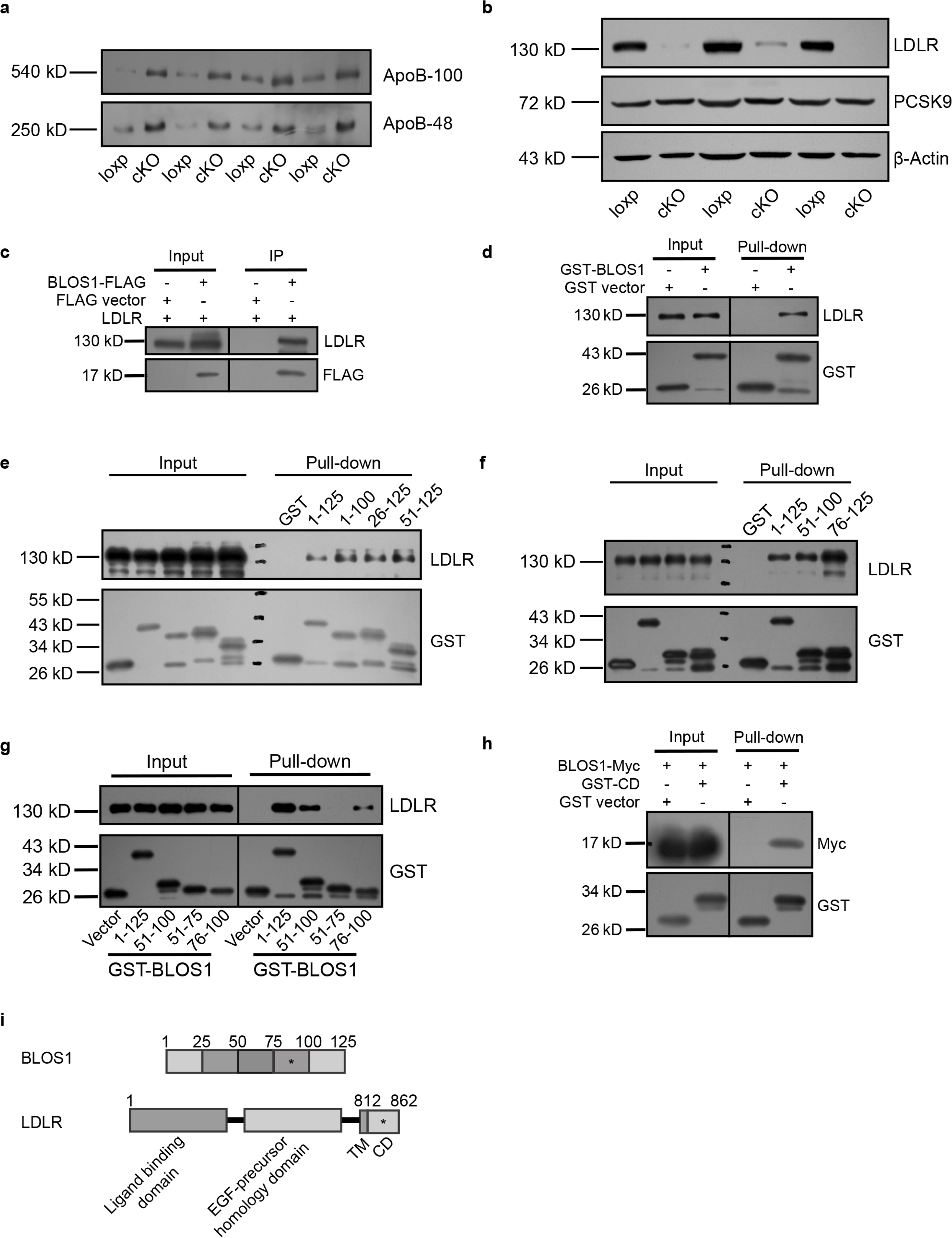
Reduced LDLR in cKO mice and LDLR interacting region mapping on BLOS1, Related to **Fig. 2**. (**a**) Immunoblot of ApoB proteins in the plasma of loxp and cKO mice. (**b**) Immunoblot of LDLR (with another LDLR antibody) and PCSK9 in loxp and cKO mice livers. (**c**) Co-immunoprecipitation (co-IP) of LDLR with co-overexpressed BLOS1-FLAG in HEK293T whole cell lysate after incubation with anti-FLAG beads, followed by immunoblotting. (**d**) Pull-down of LDLR in mouse liver lysate by GST-BLOS1 purified with glutathione sepharose beads, followed by immunoblotting. (**e**, **f**) Pull-down of LDLR with different BLOS1 truncations fused to GST. BLOS1 was truncated from both sides, and the interacting region was further narrowed with shorter truncations. (**g**) Pull-down of LDLR by different truncations of BLOS1 fused to GST, showing the interacting region on BLOS1. (**h**) Pull-down of BLOS1-Myc in HEK293T cell lysate by GST-CD (cytoplasmic domain of LDLR fused to GST). (**i**) Schematic diagram of mapping of the interacting regions on BLOS1 and LDLR, the star sign (*) represents the mapped region.

**Figure S3.**
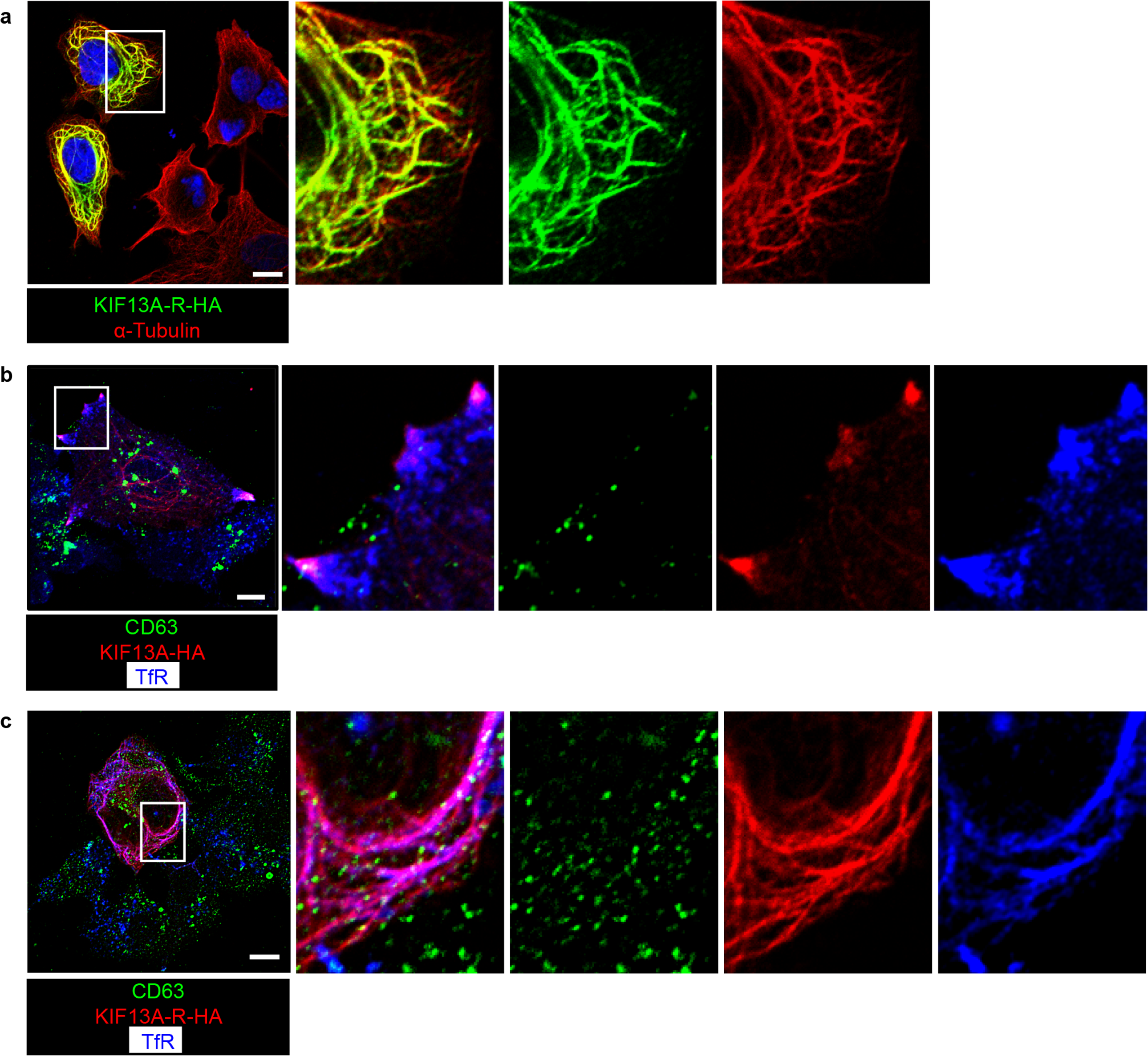
KIF13A and its rigor mutant do not affect lysosome distribution, Related to **Fig. 4**. (**a**) Representative confocal image showing KIF13A-R distributes along a proportion of microtubules (indicated by α-Tubulin). (**b**) Representative confocal image showing that overexpressed full-length KIF13A redistributes recycling endosomes (indicated by TfR) but not lysosomes/MVB (indicated by CD63) to the cell periphery. (**c**) Representative confocal image showing that KIF13A-R retains recycling endosomes (indicated by TfR) but not lysosomes/MVB (indicated by CD63) on KIF13A-R-positive microtubules. Scale bars in all pictures, 10 μm.

**Figure S4.**
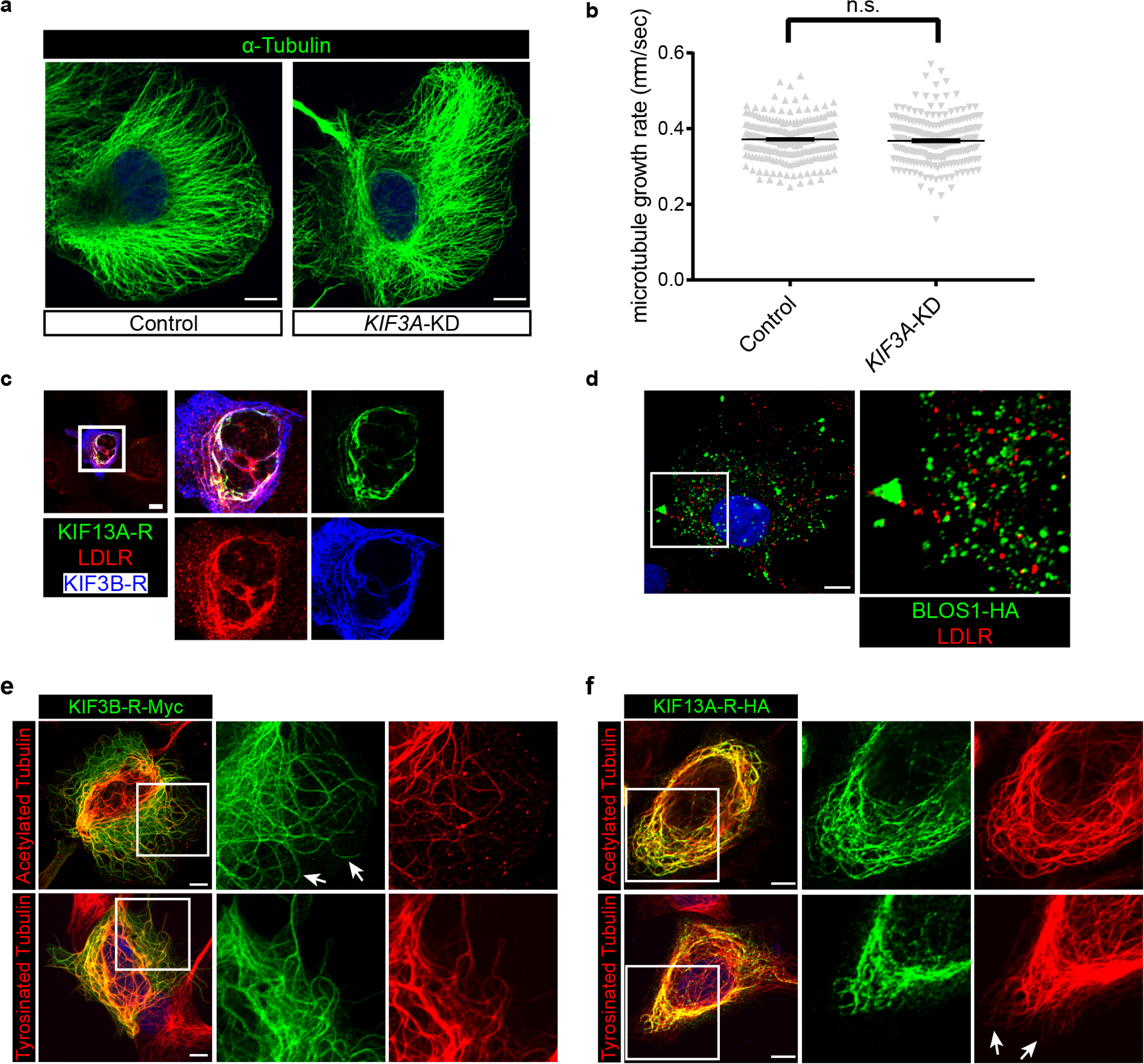
Kinesin-2 and kinesin-3 bind to the same set of microtubule tracks with acetylation tubulin, Related to **Fig. 6**. (**a**) Representative confocal image of microtubules (stained by α-Tubulin) in control and KIF3A-KD cells. (**b**) Quantification of the microtubule growth rate (indicated by the movement of EB1-GFP labeled positive tips of microtubules) in control (n = 231 tips) and KIF3A-KD (n = 224 tips) cells. Date are presented as Mean ± SEM. Two-tailed t test; n.s., not significant. (**c**) Immunofluorescence staining of LDLR after co-expression of KIF13A-R-HA and KIF3B-R-Myc. Merged and single labeling images of magnified insets of boxed area are shown on right. (**d**) Immunostaining of LDLR showing the distribution of LDLR after the co-overexpression of KIF3A-FLAG, KIF3B-Myc and BLOS-HA using multiple expression cassette-containing plasmids. (**e**) Representative confocal image showing the distribution of KIF3B-R on acetylated and tyrosinated microtubules. Arrows indicate the KIF3B-R-positive microtubules without acetylated Tubulin. (**f**) Representative confocal images showing the distribution of KIF13A-R on acetylated and tyrosinated microtubules. Arrows indicate the tyrosinated microtubules without KIF13A-R binding. Scale bars in all pictures, 10 μm.

## Supplementary Video Legends

**Video 1 (related to Fig. 4):** Live-cell imaging of Hep G2 cells transfected with KIF13A-R-Scarlet and RE marker RAB11A-GFP. Cells were recorded every 1 s for 120 s. Panels on the right show magnified views of the boxed region. Note those almost static REs on KIF13A-R tubules during the imaging period.

**Video 2 (related to Fig. 5):** Live-cell imaging of Hep G2 cells transfected with KIF3B-R-Scarlet and (or without) RE marker RAB11A-GFP. Cells were recorded every 1 s for 120 s. Panels on the right show magnified views of REs in boxed region. Note the active movement of REs in cells without KIF3B-R expression (top right) and impaired RE trafficking in KIF3B-R expressing cells (bottom right).

**Video 3 (related to Fig. 6):** Live-cell imaging of REs in control and *KIF3A*-KD cells. Control (left) and *KIF3A*-KD (right) cells expressing RAB11A-GFP (grayscale) were recorded every 1 s for 120 s. Representative RE vesicles are outlined by circles with distinct color and trajectories are showed in a backward (fast) manner. Arrows on the left indicate pausing points of REs in control cells. Note the reversed moving polarity of circled REs in *KIF3A*-KD cells after paused anterograde transport.

**Video 4 (related to Fig. 6):** Live-cell imaging of REs in *BLOC1S1*-KD cells. Images were acquired and processed as described in**Video 3**.

**Video 5 (related to Fig. 6):** Live-cell imaging of KIF13A tubules in control and *KIF3A*-KD cells. Control (left) and *KIF3A*-KD (right) cells expressing KIF13A-GFP were recorded every 1 s for 120 s. Boxed regions in control and *KIF3A*-KD cells are magnified and showed in the latter part of the movie. Filled arrowheads indicate the tips of KIF13A tubules in control cells that could successfully reach the cell periphery, and the open arrowheads indicate the tip of KIF13A tubules in *KIF3A*-KD cells that retracted after they encounter other KIF13A tubules.

**Video 6 (related to Fig. 6):** Live-cell imaging of microtubule dynamics in control (left) and *KIF3A*-KD (right) cells. The (+) tips of microtubules were labeled with EB1-GFP, and the movement of these EB1-GFP tips were recorded every 1 s for 120 s.

**Video 7 (related to Fig. 7):** Live-cell imaging of recycling endosomes (indicated by RAB11A-GFP) moving on microtubule tracks (stained by Tubulin Tracker Deep Red probe) showing the passage of recycling endosomes after pausing at specific microtubule intersections. Arrows indicate intersections where recycling endosomes (marked with magenta circles) pause and then go through. Images were acquired with Zeiss LSM 880 using the Airyscan module with line mode, and the time interval between every two frames was 0.32 second.

**Video 8 (related to Fig. 7):** Live-cell imaging of recycling endosomes (indicated by RAB11A-GFP) moving on microtubule tracks (stained by Tubulin Tracker Deep Red probe) showing the reversed movement of recycling endosomes at specific microtubule intersections. Arrows indicate intersections where recycling endosomes (marked with magenta circles) pause and then reversed their moving directions. Images were acquired with Zeiss LSM 880 using the Airyscan module with line mode, and the time interval between every two frames was 0.32 second.

**Video 9 (related to Fig. 8):** Live-cell imaging of BLOS1-GFP puncta showing the frequent z-position change of BLOS1 puncta during recording period. Two typical BLOS1 puncta are marked by dashed circles, and their intensity profiles are plotted on the right. Arrows indicate the time points at which BLOS1 puncta are in focus and show higher intensity. The time interval between every two frames was 0.58 second.

**Table S1.** Key resources.

| Reagent or resource | Source | Identifier |
| --- | --- | --- |
| Antibodies |  |  |
| Mouse monoclonal anti-alpha Tubulin (clone DM1A) | Abcam | Cat#ab7291;<br>RRID:AB_2241126 |
| Rabbit polyclonal anti-alpha Tubulin | Abcam | Cat#ab18251,<br>RRID:AB_2210057 |
| Rabbit polyclonal anti-Transferrin Receptor | Abcam | Cat#ab84036,<br>RRID:AB_10673794 |
| Rabbit polyclonal anti-LDL Receptor | Abcam | Cat#ab30532,<br>RRID:AB_881272 |
| Rabbit monoclonal anti-LDL Receptor (clone EP1553Y) | Abcam | Cat#ab52818,<br>RRID:AB_881213 |
| Goat polyclonal anti-HA tag antibody | Abcam | Cat#ab9134,<br>RRID:AB_307035 |
| Rat monoclonal anti-mouse IgG for IP (HRP) | Abcam | Cat#ab131368, N/A |
| Donkey anti-rabbit Alexa Fluor 405 | Abcam | Cat#ab175649,<br>AB_2715515 |
| Mouse monoclonal anti-EEA1 (clone 14) | BD | Cat#610457,<br>RRID:AB_397830 |
| Mouse monoclonal anti-Cytochrome C (clone 6H2.B4) | BD | Cat#556432,<br>RRID:AB_396416 |
| Mouse monoclonal anti-CD63 (clone H5C6) | BD | Cat#556019,<br>RRID:AB_396297 |
| Goat polyclonal anti-mouse LDL Receptor | R&D | Cat#AF2255,<br>RRID:AB_355203 |
| Goat polyclonal anti-human LDL Receptor | R&D | Cat#AF2148,<br>RRID:AB_2135126 |
| Mouse monoclonal anti-beta Actin (clone AC-15) | Sigma-Aldrich | Cat#A5441,<br>RRID:AB_476744 |
| Mouse monoclonal anti-Acetylated Tubulin antibody (clone 6-11B-1) | Sigma-Aldrich | Cat#T7451,<br>RRID:AB_609894 |
| Mouse monoclonal anti-FLAG tag antibody (clone M2) | Sigma-Aldrich | Cat#F3165,<br>RRID:AB_259529 |
| Rabbit polyclonal anti-FLAG tag antibody | Sigma-Aldrich | Cat#F7425,<br>RRID:AB_439687 |
| Rat monoclonal anti-Tyrosinated Tubulin antibody (clone YL1/2) | Millipore | Cat#MAB1864,<br>RRID:AB_2210391 |
| Rabbit monoclonal anti-KIF3A antibody (clone D7G3) | Cell Signaling Technology | Cat#8507,<br>RRID:AB_11141049 |
| Rabbit polyclonal anti-KIF13A antibody | Bethyl Laboratories | Cat#A301-077A,<br>RRID:AB_873053 |
| Rabbit polyclonal anti-Myc tag antibody | MBL | Cat#562,<br>RRID:AB_591105 |
| Mouse monoclonal anti-Myc tag antibody (clone Myc3) | MBL | Cat#M192-3S,<br>AB_11161202 |
| Mouse monoclonal anti-HA tag antibody (clone F-7) | Santa Cruz | Cat#sc-7392,<br>RRID:AB_627809 |
| Mouse monoclonal anti-GST antibody (clone B-14) | Santa Cruz | Cat#sc-138,<br>RRID:AB_627677 |
| Rabbit polyclonal anti-Pallidin antibody | Proteintech | Cat#10891-2-AP,<br>RRID:AB_2164174 |
| Rabbit polyclonal anti-Dysbindin antibody | Prepared in our lab<br>(Wang et al., 2014) | N/A |
| Donkey anti-mouse Alexa Fluor 488 | ThermoFisher | Cat#A-21202,<br>RRID:AB_141607 |
| Donkey anti-mouse Alexa Fluor 594 | ThermoFisher | Cat#A-21203,<br>RRID:AB_2535789 |
| Donkey anti-Rabbit Alexa Fluor 488 | ThermoFisher | Cat#A-21206,<br>RRID:AB_2535792 |
| Donkey anti-Rabbit Alexa Fluor 594 | ThermoFisher | Cat#A-21207,<br>RRID:AB_141637 |
| Donkey anti-Goat Alexa Fluor 488 | ThermoFisher | Cat#A-11055,<br>RRID:AB_2534102 |
| Donkey anti-Goat Alexa Fluor 594 | ThermoFisher | Cat#A-11058,<br>RRID:AB_2534105 |
| Bacterial and Virus Strains |  |  |
| DH5α chemically component cells | Vazyme | Cat#C502 |
| BL21 (DE3) chemically component cells | Vazyme | Cat#C504 |
| Chemicals, Peptides, and Recombinant Proteins |  |  |
| Minimal Essential Medium (MEM) | GE Healthcare | Cat#SH30024.01 |
| Glutathione Sepharose 4B resin | GE Healthcare | Cat#17075601 |
| Puromycin | InvivoGen | Cat#ant-pr-1 |
| Leupeptin | Sigma-Aldrich | Cat#L5793 |
| Collagen, Type I | Sigma-Aldrich | Cat#C3867 |
| Collagenase, Type II | Sigma-Aldrich | Cat#C5138 |
| Oil Red O | Sigma-Aldrich | Cat# O9755 |
| Sudan Black B | Sigma-Aldrich | Cat#199664 |
| 1,1'-Dioctadecyl-3,3,3',3'-tetramethylindocarbocyanine perchlorate (Dil) | Sigma-Aldrich | Cat#42364 |
| Anti-FLAG M2 affinity gel | Sigma-Aldrich | Cat#A2220 |
| Phalloidin Alexa Fluor 594 | ThermoFisher | Cat#A12381 |
| Prolong Gold Antifade Mountant | ThermoFisher | Cat#P36935 |
| Lipofectamine 3000 | ThermoFisher | Cat#L3000015 |
| Sodium pyruvate | ThermoFisher | Cat#11360070 |
| MEM Non-Essential Amino Acids | ThermoFisher | Cat#11140050 |
| Tubulin Tracker Deep Red | ThermoFisher | Cat# T34076 |
| jetPEI-Hepatocyte | Polyplus | Cat#102-05N |
| Critical Commercial Assays |  |  |
| iScript cDNA Synthesis Kit | BIO-RAD | Cat#1708891 |
| RNeasy Mini Kit | QIAGEN | Cat#74104 |
| ClonExpress II One Step Cloning Kit | Vazyme | Cat#C112 |
| Experimental Models: Cell Lines |  |  |
| Human: Hep G2 | Cell bank of Chinese Academy of Sciences (Shanghai, China) | Cat#TCHu72, RRID:CVCL_0027 |
| Mouse primary cultured hepatocytes | This paper | N/A |
| Experimental Models: Organisms/Strains |  |  |
| Mouse: loxp | Prepared in our lab (Zhang et al., 2014) | N/A |
| Mouse: cKO | This paper | N/A |
| Mouse: *pa*: B6.Cg-Bloc1s6<sup>pa</sup>/J | The Jackson Laboratory | JAX: 000024, RRID:IMSR_JAX:000024 |
| Mouse: *Kxd1*-KO | Prepared in our lab (Yang et al., 2012) | N/A |
| Mouse: Alb-Cre: B6.Cg-Speer6-ps1<sup>Tg(Alb-cre)21Mgn</sup>/J | Model animal research center of Nanjing university | Derived from The Jackson Laboratory, RRID:IMSR_JAX:003574 |
| Oligonucleotides |  |  |
| See **Table S2** | N/A | N/A |
| Recombinant DNA |  |  |
| BLOS1-GFP-C2 | This paper | N/A |
| BLOS1-GFP-N2 | This paper | N/A |
| BLOS1-FLAG | This paper | N/A |
| BLOS1-Myc | This paper | N/A |
| BLOS1-HA | This paper | N/A |
| GST-BLOS1 | This paper | N/A |
| RAB11A-GFP-C2 | This paper | N/A |
| RAB11A-Scarlet-C2 | This paper | N/A |
| KIF13A-FLAG | This paper | N/A |
| KIF13A-GFP-N2 | This paper | N/A |
| KIF13A-ST-GFP-N2 | This paper | N/A |
| KIF13A-HA | This paper | N/A |
| KIF13A-R-HA | This paper | N/A |
| KIF13A-R-Scarlet-N2 | This paper | N/A |
| KIF3B-Myc | This paper | N/A |
| KIF3B-R-Myc | This paper | N/A |
| KIF3B-R-Scarlet-N2 | This paper | N/A |
| KIF3A-FLAG | This paper | N/A |
| KIF3A-R-FLAG | This paper | N/A |
| KIF3C-HA | This paper | N/A |
| KIF3C-R-HA | This paper | N/A |
| KAP3-HA | This paper | N/A |
| KIF5B-R-Myc | This paper | N/A |
| KIF16B-FLAG | This paper | N/A |
| KIF3A-FLAG-KIF3B-Myc-BLOS1-HA | This paper | N/A |
| Software and Algorithms |  |  |
| Fiji | http://fiji.sc/<br>(Schindelin et al., 2012) | Version 2.0.0-rc-69/1.52n,<br>RRID:SCR_002285 |
| Other |  |  |
| μ-Dish 35 mm, high Glass Bottom | *ibid* | Cat#81158 |

N/A, not applicable.

**Table S2.**
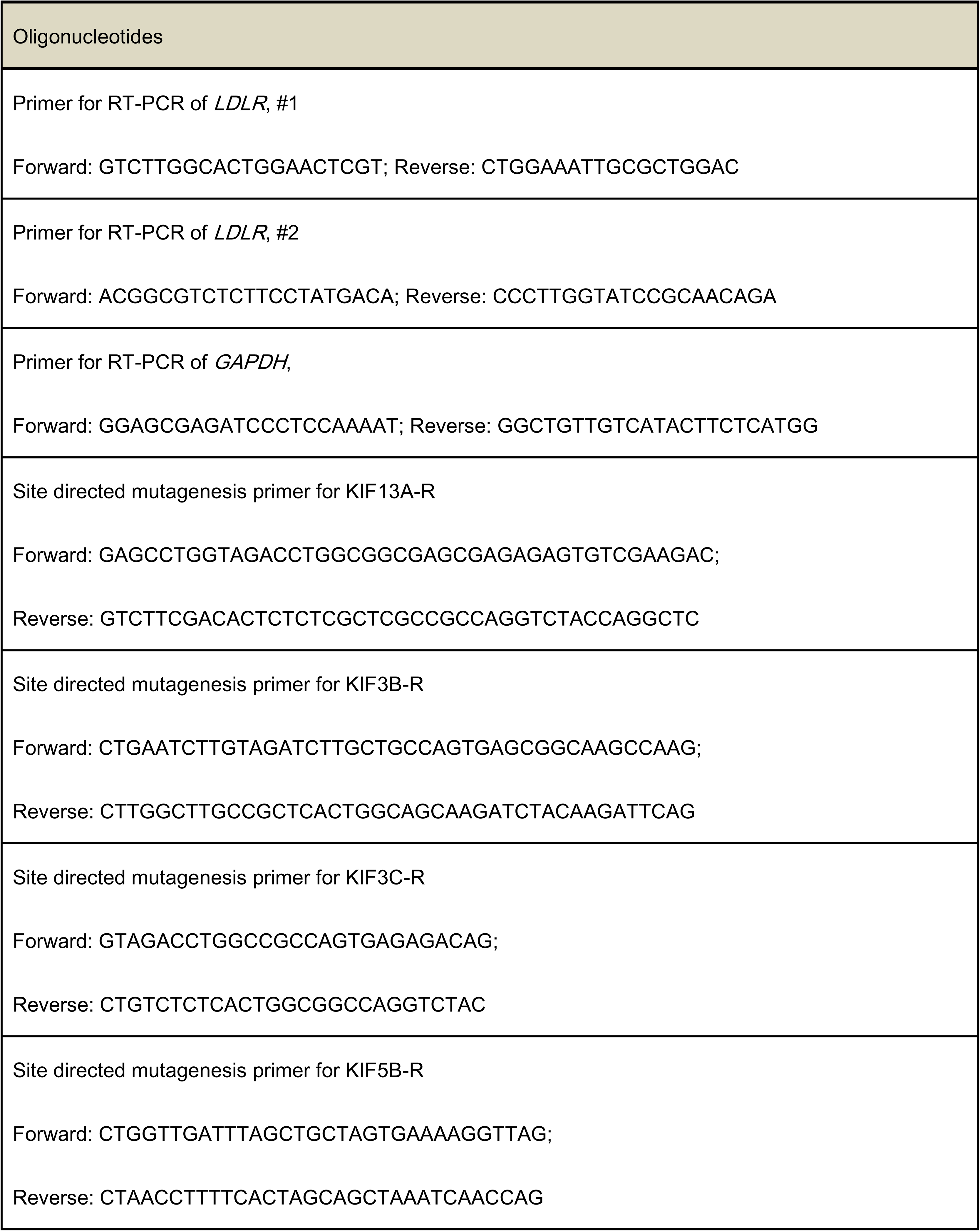
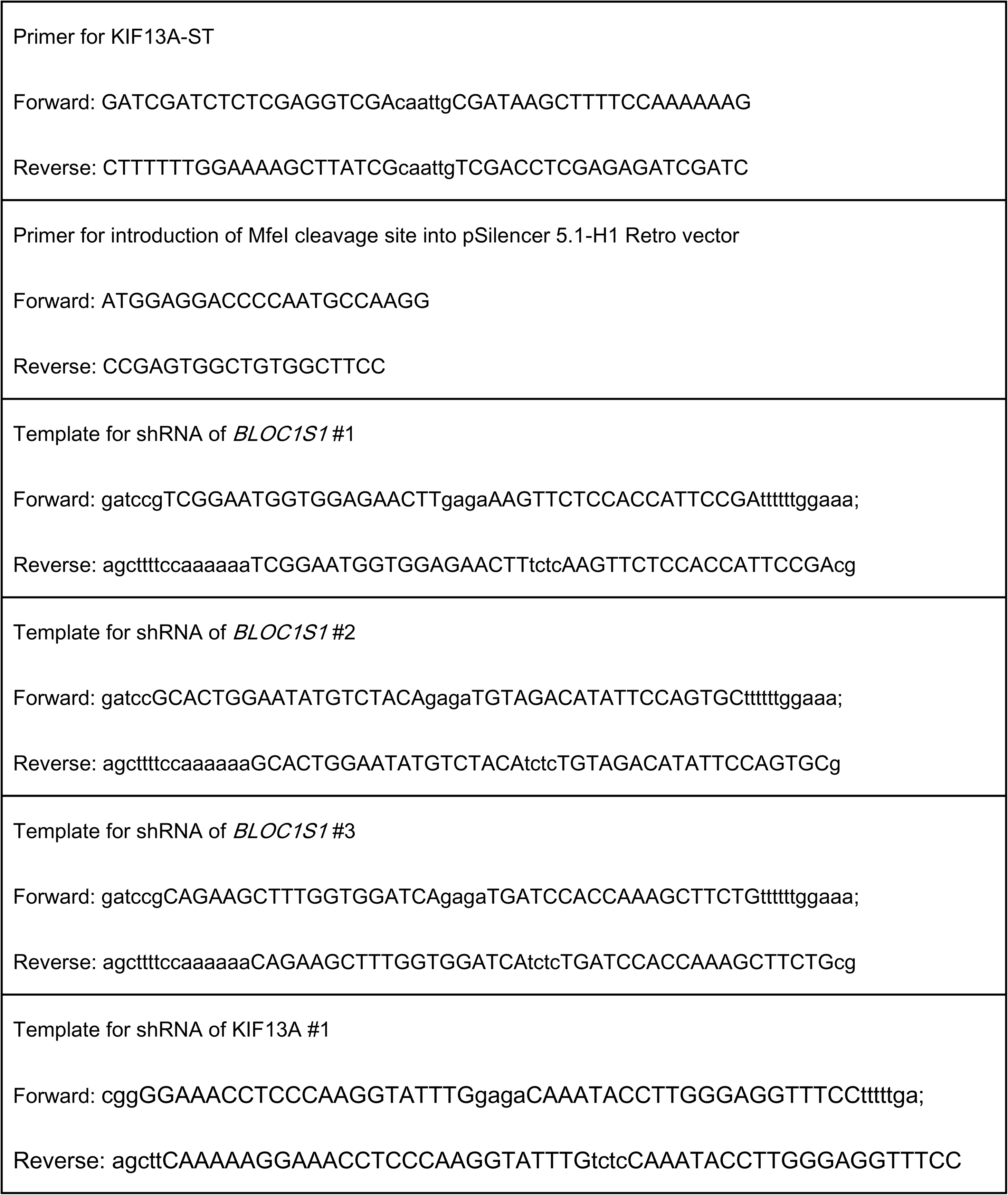
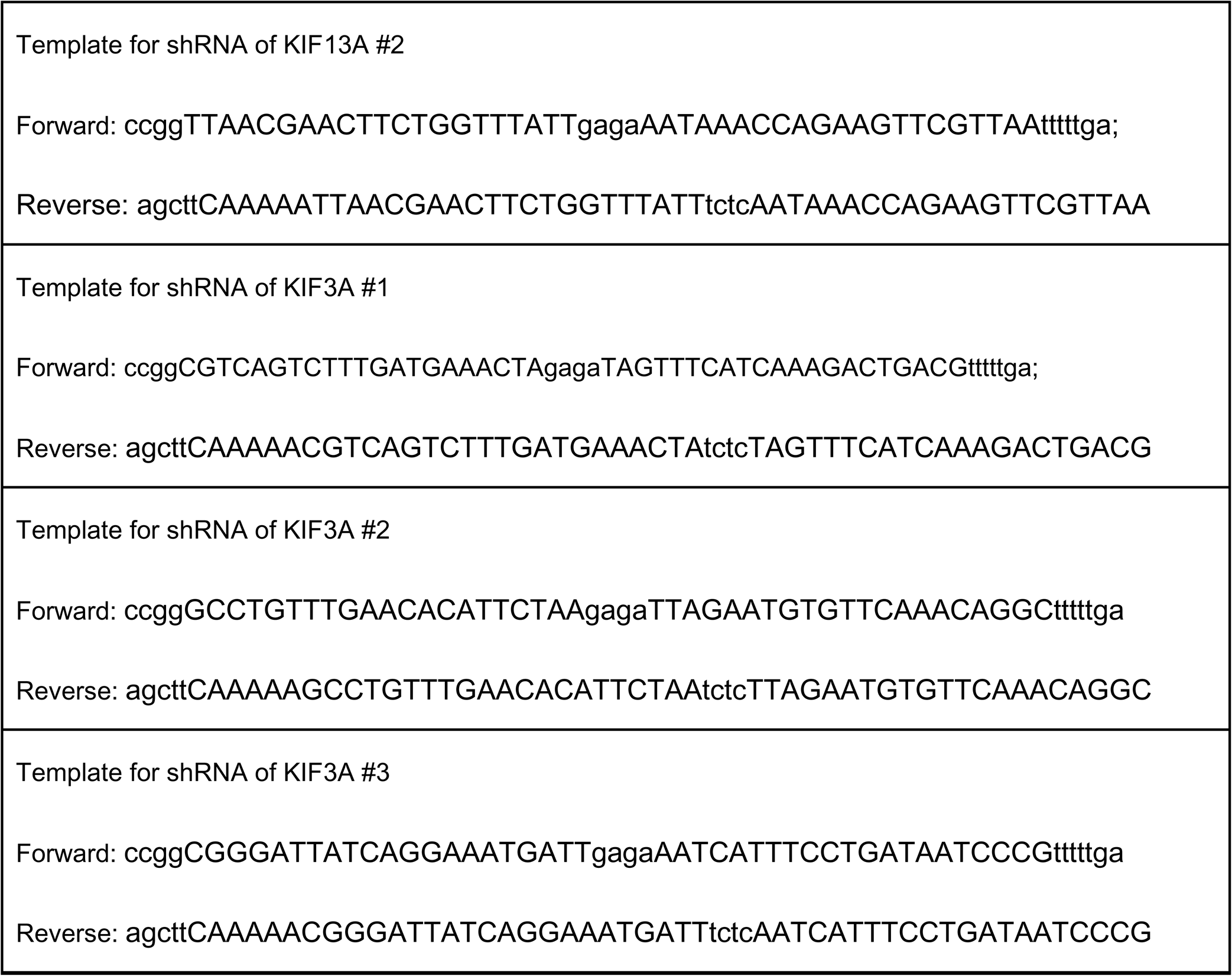
Oligonucleotides used in this study.

